# INDETERMINATE DOMAIN-DELLA protein interactions orchestrate gibberellin-mediated cell elongation in wheat and barley

**DOI:** 10.1101/2025.04.17.649374

**Authors:** Patrycja Sokolowska, Matthias Jost, Wolfram Buss, Brett Ford, Peter Michael Chandler, Wolfgang Spielmeyer, Andy Phillips, Alison K. Huttly, Danuše Tarkowská, Rocío Alarcón-Reverte, Suzanne J. Clark, Stephen Pearce, Peter Hedden, Stephen G. Thomas

## Abstract

DELLA proteins, members of the GRAS-domain family of transcriptional regulators, are critical for plant growth and development. They modulate transcription indirectly via interactions with hundreds of transcription factors. The phytohormone gibberellin (GA) triggers DELLA degradation, providing a mechanism by which plants can integrate developmental and environmental signals to regulate gene expression and optimize growth responses.

In agriculture, DELLA mutations have been instrumental in improving crop performance. Most modern wheat (*Triticum aestivum* L.) varieties carry *Rht-B1b* or *Rht-D1b* alleles that encode DELLA proteins resistant to GA-mediated degradation, resulting in constitutive partial suppression of stem growth, a semi-dwarf stature and lodging resistance. However, these alleles also reduce nitrogen use efficiency and early vigour, limiting their utility in some environments. Understanding how DELLA proteins regulate growth and development is, therefore, critical for refining breeding strategies.

In this study, we identified the orthologous C2H2 zinc-finger transcription factors *INDETERMINATE DOMAIN 5* (*IDD5*) in wheat and *SEMI-DWARF 3* (*SDW3*) in barley (*Hordeum vulgare*) as positive regulators of stem and leaf expansion. Both IDD5 and SDW3 physically interact with, and act downstream of, DELLA proteins as key components of GA-mediated growth responses. Altered expression levels of GA biosynthesis genes suggest that IDD5 helps maintain GA homeostasis in addition to growth regulation.

Loss-of-function mutations in *IDD5* and *SDW3* confer a GA-insensitive semi-dwarf phenotype comparable to that of the *Rht-D1b* ‘Green Revolution’ allele, highlighting their potential as novel dwarfing alleles for cereal improvement.

**Significance statement:** Our study identifies homologous wheat and barley transcription factors (IDD5 and SDW3) that interact with DELLA proteins to regulate plant height. Unlike conventional ‘Green Revolution’ DELLA mutations, which can reduce height but also have drawbacks such as lower nitrogen-use efficiency, these IDD genes may provide more targeted approaches to manage plant growth. By showing how IDD proteins promote stem and leaf expansion and by revealing their potential as alternative dwarfing alleles, our research opens new avenues both for fundamental research into plant growth pathways and for applications in cereal breeding. Ultimately, it could help produce crops with improved lodging resistance and fewer negative side effects than current dwarfing alleles.

## Introduction

In higher plants, DELLA proteins are critical regulators of multiple developmental processes, including cell elongation, flowering, and germination, as well as responses to biotic and abiotic stresses (1). DELLA proteins regulate these processes via transcriptional reprogramming through interactions with members of at least 15 different families of transcription factors (TFs) (2, 3). These protein-protein interactions involve conserved motifs within the C-terminal GRAS domain of DELLA proteins (3-5).

In some cases, interaction with DELLA proteins inhibits the DNA-binding activity of target TFs to suppress transcriptional activity. For example, DELLA proteins bind to and sequester PHYTOCHROME INTERACTING FACTOR (PIF4) that promotes cell elongation, and GROWTH REGULATING FACTOR 4 (GRF4) that modulates the balance between growth, N assimilation and carbon fixation (6-8). In other cases, DELLA proteins increase the transcriptional activity of bound TFs by recruiting transcriptional activation complexes via N-terminal transactivation domains (9-11).

The activity of DELLA proteins is regulated by gibberellins (GAs), a class of phytohormones that promote growth. Although 136 GA forms have been described, only a subset -such as GA_1_ and GA_4_ -are biologically active (12). The cellular levels of bioactive GAs are tightly regulated in response to developmental signals and environmental cues through the coordinated activity of different classes of GA biosynthetic and catabolic enzymes (13).

Bioactive GA binds to the receptor protein GIBBERELLIN INSENSITIVE DWARF 1 (GID1), causing a conformational change that enhances its binding affinity for DELLA proteins via conserved N-terminal domains. The resulting GA-GID1-DELLA complex is recognized by the SKP1-CULLIN-F-box (SCF)_GID2_ ubiquitin E3 ligase complex, targeting DELLA proteins for polyubiquitination and rapid 26S proteasome-mediated degradation (14). The GA-mediated degradation of DELLA proteins results in transcriptional changes that promote GA-responsive growth (3).

Mutations in DELLA genes have been exploited in agriculture to improve crop performance. Semi-dwarfing alleles of the B and D homoeologues of *RHT1*, which encode DELLA proteins in wheat (*Triticum aestivum* L.), were selected during the ‘Green Revolution’ for their positive effects on lodging resistance and assimilate partitioning that results in higher yields in some environments (15). The *Rht-B1b* and *Rht-D1b* alleles carry point mutations that introduce premature stop codons in the N-terminal DELLA domain (4, 16). Translational reinitiation at a downstream AUG codon produces an N-terminally truncated DELLA protein that cannot be bound by the GA-GID1 complex. This truncated protein is resistant to GA-mediated degradation and confers constitutive growth repression in stem tissues (17). In barley (*Hordeum vulgare*), mutations that disrupt the N-terminal DELLA domain of SLENDER1 (SLN1), the orthologous DELLA protein, confer a dominant gain-of-function semi-dwarf phenotype (18). This demonstrates the functional conservation of DELLA genes in temperate cereals.

However, in addition to reduced stem elongation, dominant, gain-of-function *Rht1* and *Sln1* alleles also confer negative pleiotropic effects, such as reduced early vigour and poor nitrogen use efficiency (NUE) that restrict their utility in some environments (19, 20). There is evidence that DELLA proteins regulate growth and development responses such as stem elongation, meristem size, and N-uptake through independent mechanisms controlled by different interacting TFs (8, 21). Therefore, a detailed understanding of the mechanisms and downstream pathways by which DELLA proteins drive different growth responses can help disentangle the effects of DELLA activity to enable more targeted strategies for cereal crop improvement.

Members of the INDETERMINATE DOMAIN (IDD) subclade of Cys2His2 (C2H2) Zinc-finger TFs interact with DELLA proteins in both monocot and dicot plant species and regulate different growth responses (2, 22). These proteins contain a conserved N-terminal ID domain consisting of four Zn-finger motifs with DNA binding activity, and a C-terminal PAM domain necessary for interaction with DELLA proteins (23-26).

The transcriptional activity of several IDD proteins is increased by binding to DELLA proteins. In *Arabidopsis*, AtIDD3 forms a protein complex with the DELLA protein GIBBERELLIC ACID INSENSITIVE (GAI) that binds to the promoter of the GA positive regulator *SCARECROW-LIKE3* (*SCL3*) to enhance its transcription (27). Similarly, the transcriptional activity of the IDD proteins ENHYDROUS (ENY, also known as AtIDD1) and GAI-ASSOCIATED FACTOR1 (GAF1, also known as AtIDD2) is increased when bound to GAI (24, 28). Unlike AtIDD3, both ENY and GAF1 have a C-terminal ERF-associated amphiphilic repression (EAR) motif that coordinates interaction with TOPLESS (TPL) proteins (28). TPL proteins recruit histone deacetylases to repress transcriptional activity (29). Therefore, one model proposes that GAF1/ENY IDD proteins act as signal-dependent TFs that form either transcriptionally repressive (via TPL) or activating (via DELLA) protein complexes, depending on cellular GA levels (27, 28).

In the current study, we demonstrate that the orthologous IDD transcription factors INDETERMINATE DOMAIN 5 (TaIDD5) in wheat and SEMI-DWARF 3 (HvSDW3) in barley physically interact with DELLA proteins and act as positive downstream regulators of GA-dependent stem and leaf expansion. The loss-of-function *idd5* and *sdw3* alleles confer a GA-insensitive semi-dwarf stature and are potential novel targets for crop improvement.

## Results

### *HvSDW3* encodes an INDETERMINATE DOMAIN (IDD) transcription factor

We previously mapped *HvSDW3* to a 5.9 Mbp region of chromosome arm 2HS in the barley genome that contains four candidate genes (Figure 1A, (30)). To determine which of these genes encodes *SDW3*, we sequenced the transcriptomes of wild-type ‘Himalaya’, M287 (a ‘Himalaya’ BC_3_ line carrying the *sdw3* allele) and seven induced mutants derived from ‘Himalaya’ that exhibit a GA-insensitive semi-dwarf growth habit and which were previously shown to be allelic to *sdw3* by complementation studies (30). Only one gene in the genome (*HORVU*.*MOREX*.*r3*.*2HG0131300*) carried SNPs in all eight sequenced mutants (Figure 1B and Figure S1). The candidate gene was located at 139,466,760 bp on chromosome arm 2HS of the Morex v3 barley genome assembly, just upstream of the outer proximal flanking marker of the previously identified physical interval (TC142185 139,469,633 bp, Table S1 (30)).

**Figure 1.**
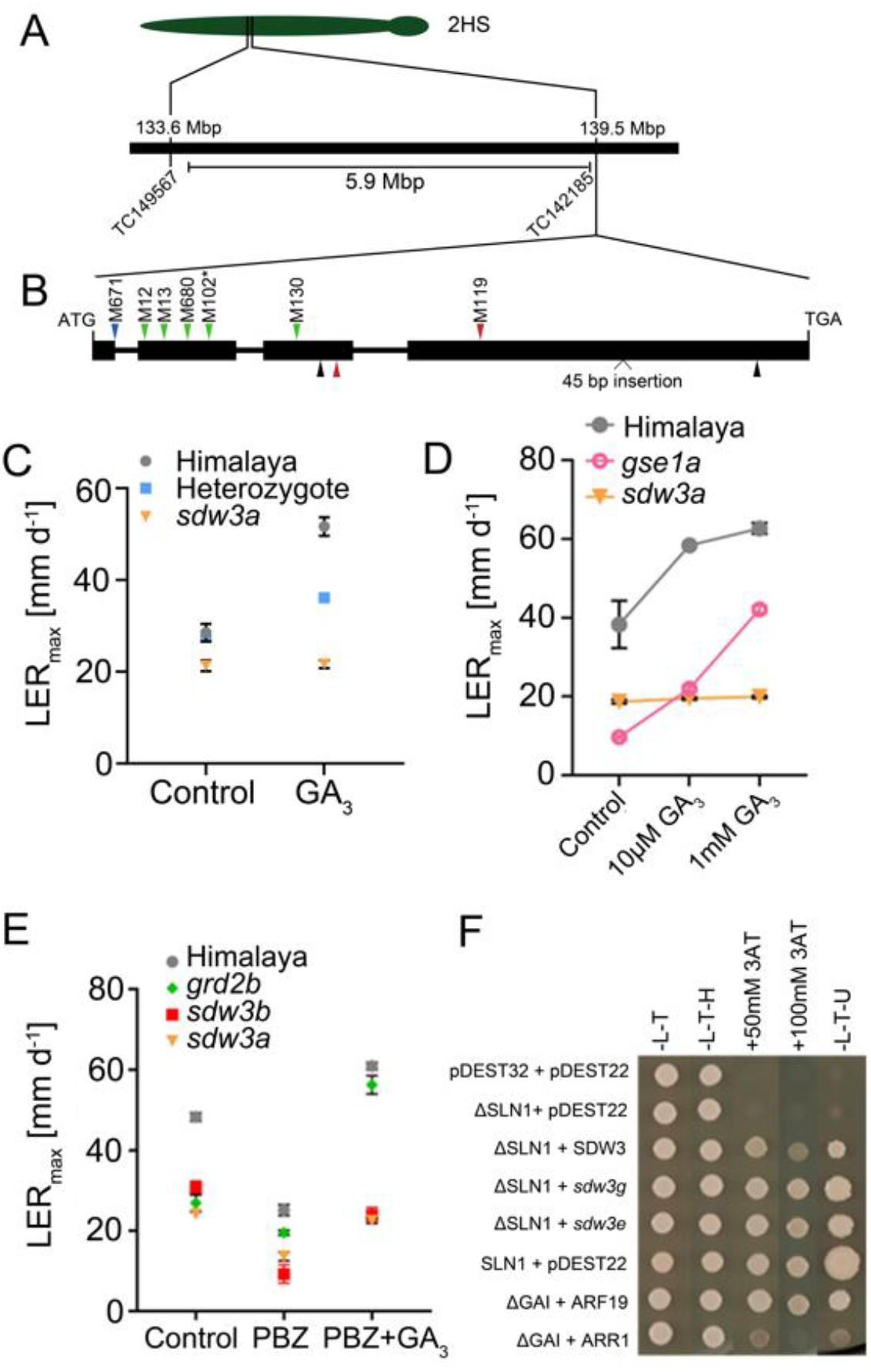
*HvSDW3* encodes an IDD transcription factor **(A)** *HvSDW3* gene structure showing previous genetic mapping delimiting a 5.9 Mbp interval in the Morex v3 barley genome assembly by placing flanking markers from (30) **(B)** Gene structure of *HvSDW3* showing the position of induced polymorphisms in ‘Himalaya’ (above the gene model) and the natural mutations in M287 (below the gene model). Different coloured arrows indicate the mutation’s functional effect (green = non-synonymous amino acid substitution, black = synonymous amino acid substitution, blue = splice site mutation, red = premature stop codon). * The mutation in M102 was also found in M107, M127 and M651 (Table 1) **(C)** Maximal rate of first leaf elongation in a population segregating for the *sdw3a* allele in response to 10 μM GA_3_ application. **(D)** Maximal rate of first leaf elongation of *sdw3a* and *gse1a* mutants in response to low (10 μM) and high (1 mM) concentrations of exogenous GA_3_. **(E)** Maximal rates of first leaf elongation of barley mutants in response to treatment with PBZ (10 μM) or PBZ plus GA_3_ (both at 10 μM). Values are means ± SEM. **(F)** Yeast-2 hybrid assays to test the interaction between HvSLN1 and HvSDW3 proteins. GAI + ARF19 and GAI + ARR1 interactions were included as strong and weak positive controls.

We confirmed the polymorphisms in *HORVU*.*MOREX*.*r3*.*2HG0131300* in the eight mutants by Sanger sequencing and identified four additional induced semi-dwarf mutant lines that also carried mutations in *HORVU*.*MOREX*.*r3*.*2HG0131300*. In total, we identified seven independent *HORVU*.*MOREX*.*r3*.*2HG0131300* alleles that are predicted to disrupt protein function (Table 1). In line M287, *HORVU*.*MOREX*.*r3*.*2HG0131300* carries four polymorphisms in the coding region, including a SNP that introduces a premature stop codon (Table 1).

**Table 1.** Induced and natural allelic variation induced in *HvSDW3*. Nucleotide and amino acid positions are based on the *SDW3* sequence in ‘Himalaya’ relative to the translation start codon. Sorting Intolerant From Tolerant (SIFT) scores for amino acid substitutions are shown in parentheses, where values less than 0.05 indicate that the mutation is predicted to have a deleterious effect on protein function. * indicates the introduction of a premature stop codon. Full sequences of all alleles are provided in additional file 1.

| Line | Allele | Nucleotide change | Effect |
| --- | --- | --- | --- |
| M671, M674 | *sdw3a* | G91A | Intron 1 splice donor site |
| M12 | *sdw3b* | C214T | S40L (0.00) |
| M13 | *sdw3c* | G277A | R61K (0.01) |
| M680 | *sdw3d* | C409T | P105L (0.00) |
| M102, M107, M127, M651 | *sdw3e* | G424A | G110E (0.00) |
| M130 | *sdw3f* | G843T | E212* |
| M5 | *sdw3g* | C1642T | L404F (0.00) |
| M287 | *sdw3(Hv287)* | A909C | E233D (0.00) |
|  |  | C1027T | Q273* |
|  |  | 45 bp insertion at 2316 bp | 15 amino acid insertion |
|  |  | G2914T | Synonymous |

In an M671 x ‘Himalaya’ F_2_ population segregating for the *sdw3a* allele, seedlings carrying the wild-type *SDW3* allele exhibited a significant increase in leaf extension in response to GA treatment (*P* < 0.001), whereas those carrying the *sdw3a* allele did not respond (*P* = 0.812, Table S2, Figure 1C). Heterozygous lines exhibited an intermediate growth response, demonstrating that mutations in *HORVU*.*MOREX*.*r3*.*2HG0131300* are associated with restrictions in cell elongation. A separate assay confirmed that barley seedlings carrying the *sdw3a* allele were insensitive to both low (10 μM) and high (1 mM) concentrations of exogenous GA_3_, in contrast to wild-type seedlings and the GA receptor mutant *gse1a* (Figure 1D, Table S3).

We next tested whether the lack of GA-responsive growth in *sdw3* mutants is due to GA insensitivity or to saturation of the GA response. Treatment with the GA biosynthesis inhibitor paclobutrazol (PBZ) reduced the rate of first leaf elongation of wild-type ‘Himalaya’, the GA biosynthesis dwarf mutant *grd2b*, and of *sdw3a* and *sdw3b* mutants (Figure 1E). Although subsequent GA_3_ application of PBZ-treated seedlings was sufficient to partially restore the rate of first leaf elongation to near control levels in *sdw3a* and *sdw3b* mutants, the effect was far less than in wild-type ‘Himalaya’ and the *grd2b* mutant, which both exhibited a much greater increase in growth response to exogenous GA_3_ (Figure 1E, Table S4). These results indicate that *sdw3* mutants are not strictly insensitive to GA_3_; rather, under normal growth conditions, one or more of the steps in the GA signalling pathway limit the response.

*HORVU*.*MOREX*.*r3*.*2HG0131300* encodes a C2H2-Zinc finger transcription factor in the IDD subfamily that is most similar to the Arabidopsis proteins ENY and GAF1 (Figure S2). Both ENY and GAF1 interact with the DELLA protein GAI (24, 28), so we tested whether HORVU.MOREX.r3.2HG0131300 interacts with the barley DELLA protein SLN1. Full-length SLN1 protein exhibits auto-activation, so we tested the interaction using an N-terminally truncated SLN1 protein (ΔSLN1) (Figure 1F). The full-length HORVU.MOREX.r3.2HG0131300 protein interacted strongly with ΔSLN1 (Figure 1F). The interaction was not affected by substitutions of conserved amino acids in a region downstream of the PAM domain (L404F in *sdw3g*) or within the ID domain (G110E in *sdw3e*), demonstrating that the reduction in height of these lines is not due to disruption of the SDW3-DELLA interaction (Figure 1F, Figure S3).

Taken together, our identification of eight independent alleles in *HORVU*.*MOREX*.*r3*.*2HG0131300* that confer reduced growth responses and the role of homologous IDD genes in GA-mediated growth responses (27, 28) led us to conclude that *HORVU*.*MOREX*.*r3*.*2HG0131300* is the causative gene for the *SDW3* locus in barley. The gene will hereafter be referred to as *HvSDW3*.

### IDD5 interacts with DELLA proteins in wheat

To identify GA signalling components in wheat, we screened an aleurone cDNA library using a ΔRHT-D1 bait protein and identified three independent clones that encode partial sequences matching the A, B and D homoeologues of *TaIDD5*, the wheat orthologue of *HvSDW3* (Figure S2). All three encoded proteins included the C-terminal PAM domain and EAR motif but lacked the N-terminal ID domain. The full-length gene sequences encode IDD proteins that share >88% amino acid identity with HvSDW3 (Figure S4).

We assembled a series of truncated IDD5-A1 proteins to test the requirement of different domains for the interaction with RHT1 (Figure 2A). We confirmed that full-length IDD5-A1 interacted strongly with ΔRHT-D1 by Y2H (Figure 2B) and validated this interaction *in vivo* using bimolecular fluorescence complementation (BiFC) and DAPI staining, which demonstrated the interaction occurs in the nucleus (Figure 2C). We assembled a truncated IDD5-A1 protein matching the three cDNA clones identified in the original Y2H screen that lack the INDETERMINATE (ID) domain and found that they exhibit stronger interaction with ΔRHT-D1 than the full-length IDD5-A1 protein (Figure 2B). This indicates that the ID domain is not necessary for the interaction between IDD5 and RHT1 and is consistent with the strong interaction between SLN1 and *sdw3e* (G110E amino acid substitution in the ID domain) (Figure 1D). IDD5 proteins lacking the PAM domain did not interact with RHT1, indicating this domain is required for the interaction (Figure 2B). The interaction between RHT1 and IDD5 proteins lacking the EAR and LDFLG motifs was weaker than with the full-length IDD5 protein, suggesting that these motifs strengthen the interaction but are not necessary (Figure 2B).

**Figure 2.**
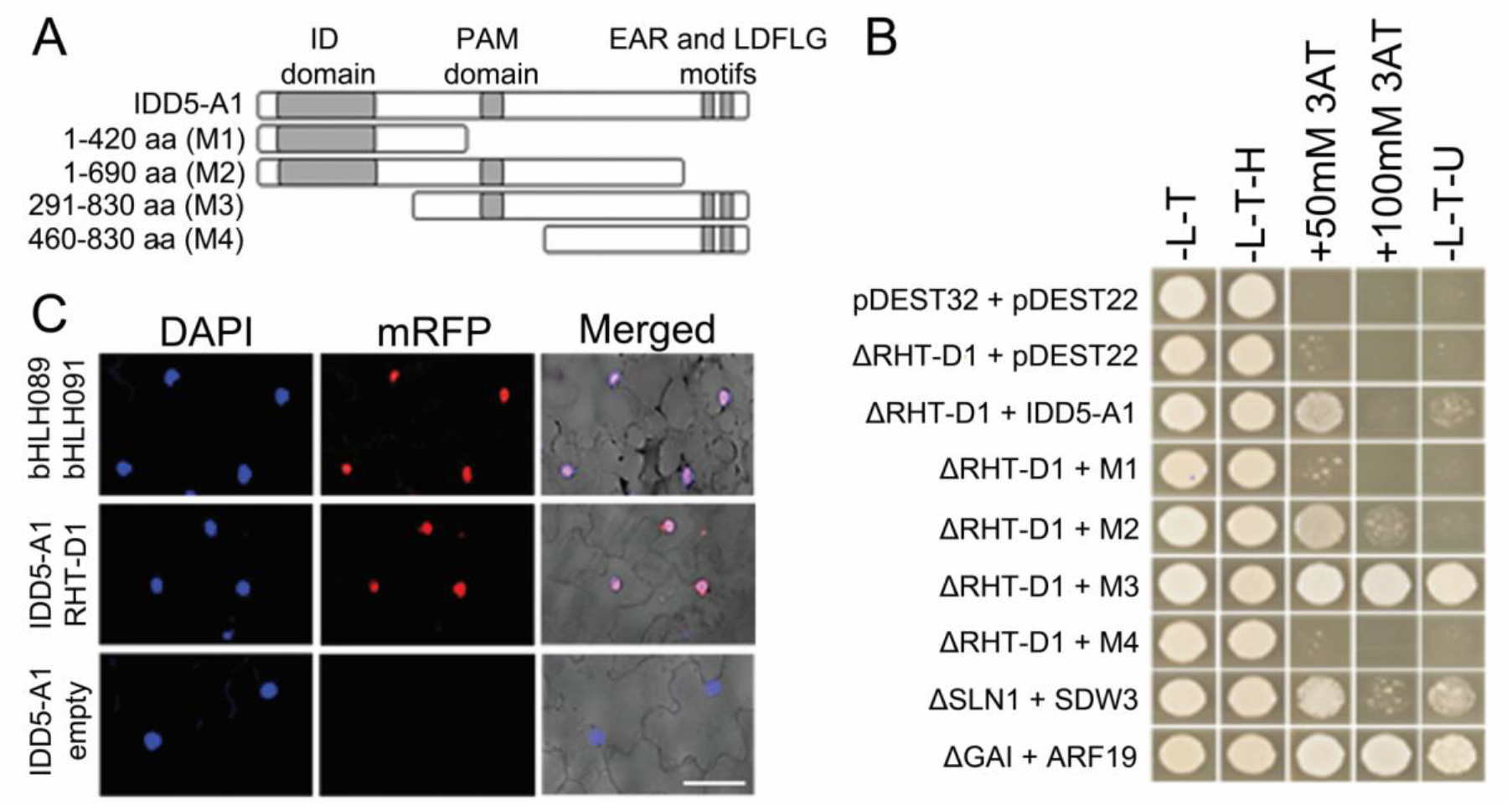
RHT1 interacts with IDD5 in wheat. **(A)** IDD5 proteins tested in Y2H assays. **(B)** Yeast-2 hybrid assays to test the interaction between RHT-D1 and IDD5-A1. GAI + ARF19 and GAI + ARR1 interactions were included as strong and weak positive controls. **(C)** *In vivo* validation of RHT-D1 – IDD5-A1 interaction using bimolecular fluorescence complementation (BiFC). bHLH089 + bHLH091 co-infiltration serves as a positive control, and TaIDD5-A1 + empty as a negative control. Scale bar = 50 µM.

Taken together, these results demonstrate that *SDW3* and *IDD5* encode orthologous IDD transcription factors that interact with DELLA proteins in barley and wheat.

### Wheat *idd5* mutants exhibit reduced GA-insensitive leaf elongation

To characterise *IDD5*, we combined three lines carrying ethyl methanesulfonate (EMS)-induced mutations in *IDD5-A1, IDD5-B1* and *IDD5-D1* to generate an *idd5* mutant carrying loss-of-function mutations in all three homoeologues (Table 2, Figure S5A). In *IDD5-A1* and *IDD5-D1*, we selected lines carrying mutations that introduce premature stop codons slightly downstream of the PAM domain, which are highly likely to encode non-functional proteins (Figure S5A). In *IDD5-B1* we selected a line carrying a mutation in the intron 1 splice donor site that corresponds to the same mutation in *sdw3a* (Figure 1B, Figure S1). The mutation in *IDD5-B1* results in a high rate of aberrant splicing producing transcripts that retain intron 1, resulting in the introduction of a premature stop codon (Figure S5B). We combined the three mutations and backcrossed three times to wild-type ‘Cadenza’ to produce a BC_3_F_3_ population segregating for each mutation. We selected lines homozygous for all three wild-type *IDD5* alleles (*IDD5*-null segregant (*IDD5*-NS) and homozygous for all three induced *idd5* mutations (*idd5*) for phenotypic characterisation.

**Table 2.** Selected EMS-induced mutations in *TaIDD5* and *RHT1* used in this study. Position in DNA relative to the translation start codon in ‘Chinese Spring’ sequences. * indicates the introduction of a premature stop codon.

| Gene | Gene ID | Position | Effect | TILLING Line |
| --- | --- | --- | --- | --- |
| *TaIDD5-A1* | *TraesCS2A02G188400* | C1909T | Q492* | CAD4-1185 |
| *TaIDD5-B1* | *TraesCS2B02G218900* | G92A | Intron 1 splice donor site | CAD4-1415 |
| *TaIDD5-D1* | *TraesCS2D02G199300* | C2046T | Q537* | CAD4-0828 |
| *TaRHT-A1* | *TraesCS4A02G271000* | G1845A | W615* | CAD4-350 |
| *TaRHT-B1* | *TraesCS4B02G043100* | G1848A | W616* | CAD4-1509 |
| *TaRHT-D1* | *TraesCS4D02G040400* | G1677A | W559* | CAD4-0587 |

Ten-day-old *idd5* seedlings were shorter than *IDD5*-NS and comparable in height to seedlings carrying the *Rht-D1b* allele (Figure 3A). The first leaf (L1) sheath was significantly shorter in *idd5* seedlings compared to *IDD5*-NS (32.4% reduction, *P* < 0.05), similar to the 36.0% reduction in *Rht-D1b* compared to wild-type ‘Cadenza’ (Figure 3B, Table S5). Whereas treatment with 100 µM GA_3_ increased the height and L1 sheath length of both ‘Cadenza’ and *IDD5*-NS seedlings, there was no effect in either *idd5* or *Rht-D1b* genotypes (Figure 3A and 3C). The L1 sheath elongation responses in both ‘Cadenza’ and *IDD5*-NS reached saturation in the range of 1 µM to 10 µM GA_3_ (Figure 3C, Table S6). By contrast, neither *idd5* nor *Rht-D1b* mutants responded to any concentration of GA_3_ (Figure 3C). These results were consistent in L1 blade elongation assays (Figure S6, Table S7). L1 sheath abaxial cell lengths were significantly different between ‘Cadenza’ and *idd5* genotypes (*P* < 0.0001) and two-way ANOVA showed a significant interaction between genotype and GA treatment (*P* = 0.18) (Figure 3D, Table S8). The effects on L1 leaf sheath length in this experiment were consistent (Figure S7, Table S9). Taken together, these results demonstrate that *idd5* mutants exhibit reduced cell elongation in leaf tissues and are insensitive to GA.

**Figure 3.**
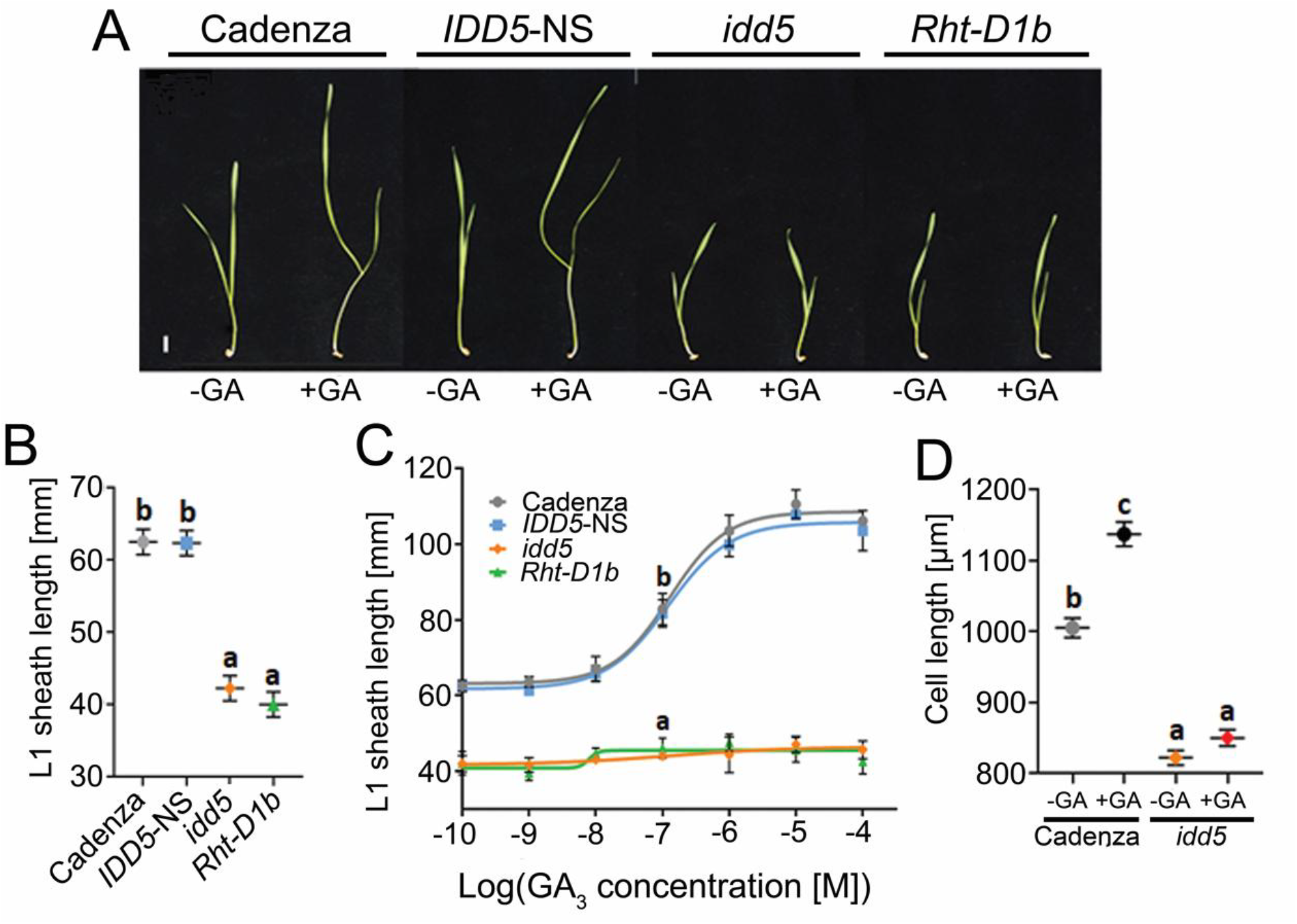
Wheat *idd5* mutants exhibit reduced cell elongation and are GA-insensitive. **(A)** The phenotype of 10-day-old seedlings of ‘Cadenza’, *Rht-D1b, IDD5*-NS and *idd5* mutants. Seedlings were either untreated (-) or sprayed with 100 µM GA_3_ every two days. **(B)** L1 sheath length in 10-day-old seedlings. Data were analysed using one-way general ANOVA followed by Tukey’s HSD test. Different letters indicate significant differences between genotypes **(C)** GA dose-response curves of L1 sheath length of seedlings 14 days after germination. Data were analysed by one-way ANOVA. **(D)** L1 sheath abaxial epidermis cell lengths of seven-day-old ‘Cadenza’ and *idd5* seedlings without (-GA) and after 100 µM GA_3_ treatment (GA). Data were analysed using unbalanced ANOVA followed by Fisher’s unprotected LSD test. All values are means ± SEM.

### IDD5 regulates GA homeostasis in the elongating leaf sheath

We quantified GA levels in the elongating leaf sheath and found that all genotypes showed higher 13-hydroxy GA levels than non-13-hydroxy GAs, consistent with previous findings that the early 13-hydroxylation pathway predominates in wheat vegetative tissues (Table S10). *Rht-D1b* and *idd5* mutants showed similar GA profiles, with both accumulating significantly higher levels of bioactive GA_1_ (2.7-fold and 2.0-fold, respectively, *P* < 0.001) and lower levels of the GA20OX substrates GA_44_ and GA_19_ than their wild-type segregants (Figure 4A). These GA profiles are consistent with increased GA 20-oxidase activity in both *idd5* and *Rht-D1b* lines. By contrast, only the *idd5* mutant exhibited significantly lower levels of GA_20_, the GA 3-oxidase substrate, consistent with increased GA 3-oxidase activity in this line (Figure 4A).

**Figure 4.**
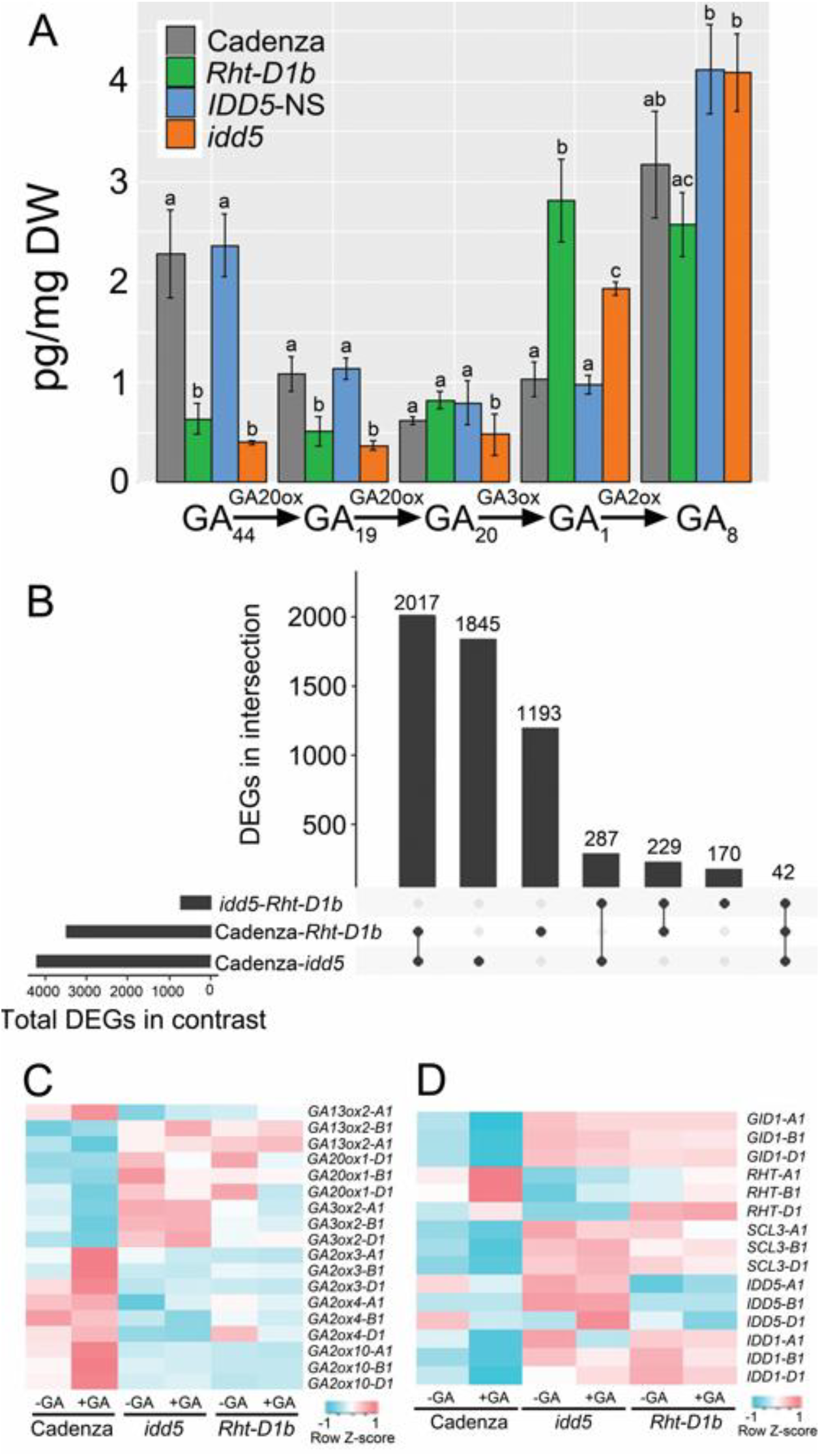
*IDD5* regulates GA levels in the wheat leaf sheath. **(A)** The levels of selected GAs in the leaf sheaths of seven-day-old seedlings of ‘Cadenza’, *IDD5*-NS, *idd5* and *Rht-D1b*. Data were analysed using one-way ANOVA and Tukey’s HSD test. Different letters indicate significant differences between genotypes (*P* < 0.05). **(B)** UpSet plot showing the number of DEGs in selected pairwise comparisons (*P*_*adj*_ < 0.01) and the subset of those genes that are common to different contrasts. Relative expression levels of selected GA biosynthesis **(C)** and GA signalling **(D)** genes in different genotypes.

We sequenced the transcriptomes of ‘Cadenza’, *Rht-D1b* and *idd5* seedlings treated with water or GA_3_ (Figure S8). In wild-type ‘Cadenza’, 364 differentially expressed genes (DEGs) were detected following GA application (Additional file 2). By contrast, the GA-insensitive *idd5* and *Rht-D1b* mutants showed only nine and one DEG, respectively (Additional file 2). Comparing *idd5* and *Rht-D1b* lines revealed 728 DEGs in water-treated seedlings (Figure 4B) and 1,302 DEGs in GA-treated seedlings (Figure S8). There were approximately five times as many DEGs between ‘Cadenza’ and either mutant (Figure 4B and 4C) with significant overlap between transcriptomes; 33.6% of genes differentially expressed between ‘Cadenza’ and *idd5* are also differentially expressed between ‘Cadenza’ and *Rht-D1b* (Figure 4B). Taken together, these results show that *idd5* and *Rht-D1b* share similar GA profiles and transcriptomes, suggesting both mutated genes act in the same genetic pathway.

In ‘Cadenza’, GA treatment reduced the expression of the GA biosynthetic genes *GA3OX2* and *GA20OX1* and increased the expression of the GA catabolic genes *GA2OX3, GA2OX4* and *GA2OX10*, consistent with their feedback regulation in maintaining GA homeostasis (Figure 4C). By contrast, *GA3OX2* and *GA20OX1* transcript levels were higher in both *idd5* and *Rht-D1b* than in ‘Cadenza’ and were unaffected by GA treatment (Figure 4C). These results are consistent with elevated GA_1_ levels and the reduced levels of the GA20OX1 substrates GA_44_ and GA_19_ (Figure 4A). Similarly, *GA2OX3*, *GA2OX4* and *GA2OX10* transcript levels were reduced in *idd5* and *Rht-D1b* mutants and did not increase after GA treatment, suggesting a disruption in GA homeostasis.

In the GA signalling pathway, both the GA receptor *GID1* and *SCL3* were upregulated in *idd5* and *Rht-D1b* compared to ‘Cadenza’ (Figure 4D), while *Rht-1* genes were downregulated in *idd5* (Figure 4D).

### TaIDD5 and HvSDW3 are epistatic to DELLA

To test the epistatic interactions between *IDD5* and *RHT1*, we developed an *rht-1* loss-of-function mutant by combining three lines carrying induced mutations introducing premature stop codons in the C-terminal SAW domain of each *RHT1* homoeologue (Table 2). The resulting *rht-1* mutant is sterile and exhibits a low-tillering, spindly growth habit resembling loss-of-function DELLA mutants in other species (Figures 5A and 5B). We crossed *rht-1* with *idd5* to generate a sextuple *idd5*/*rht-1* mutant. In the *idd5* background, the spindly, low-tillering phenotype of *rht-1* was rescued, and the height of the *idd5*/*rht-1* mutant was not significantly different from *idd5* (*P* > 0.05, Figure 5A and 5B, Table S11). Despite rescuing the height phenotype, both the *rht-1* and *idd5*/*rht-1* mutants remained infertile.

**Figure 5.**
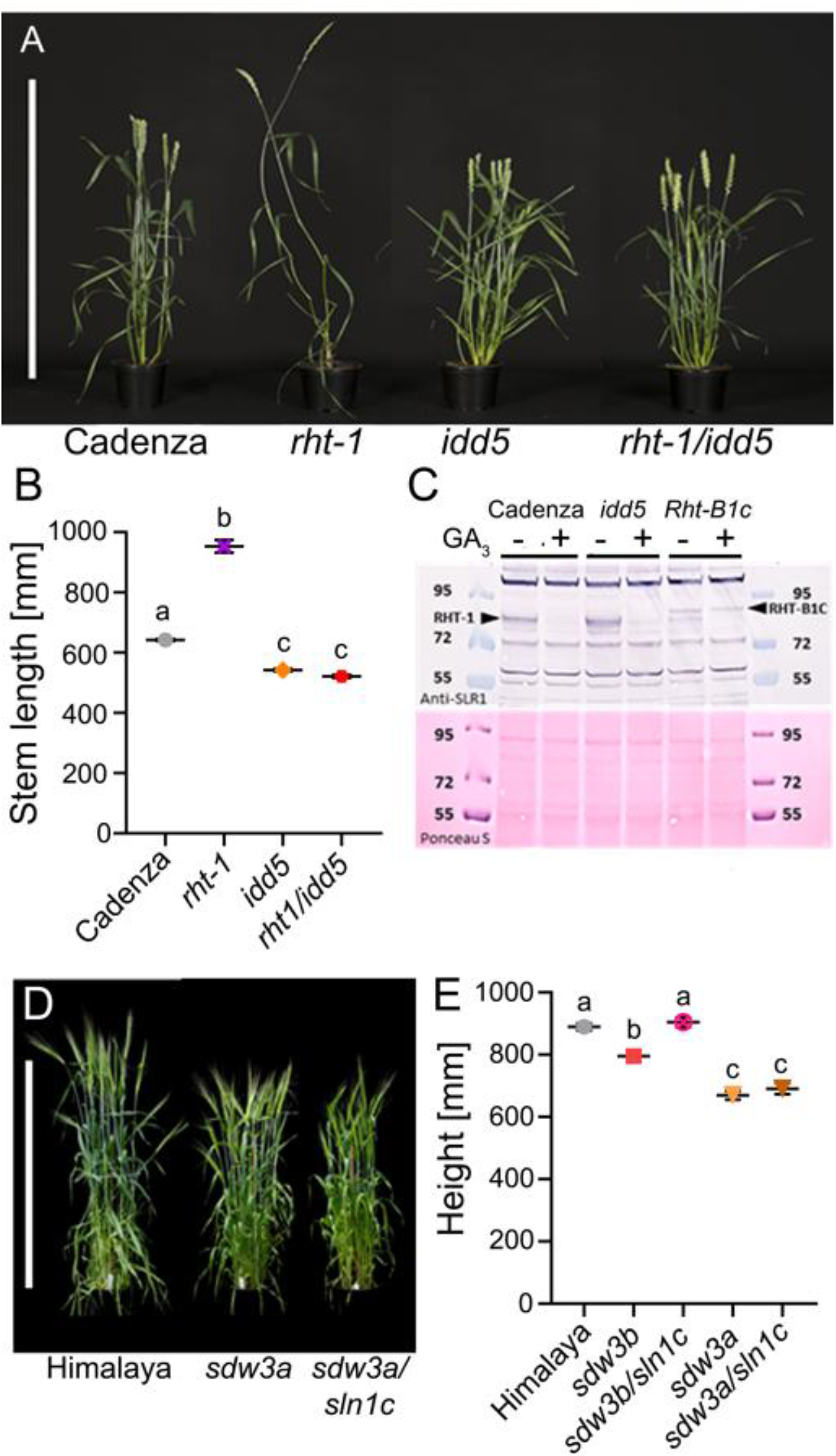
*IDD5* and *SDW3* act downstream of DELLA genes. **(A)** Whole plant phenotypes of *rht1, idd5* and *rht1/idd5* mutants at maturity. **(B)** Mean stem length (mm) ± SEM at maturity. **(C)** Detection of RHT1 proteins by Western blot in ‘Cadenza’, *idd5* and *Rht-B1c* immature spike protein extracts. Protein extracts were untreated or treated with GA_3_. RHT-1 and RHT-B1C proteins are indicated. **(D)** Whole plant phenotypes of *sdw3a* and *sdw3a/sln1c* mutant barley plants at maturity. **(E)** Total plant height at maturity. Data was analysed with one-way ANOVA and Tukey’s post-hoc HSD test. Different letters indicate significant differences between genotypes. Bar = 90 cm.

In wild-type ‘Cadenza’, native RHT-1 proteins in immature spike extracts are degraded in response to GA application (Figure 5C). By contrast, in the GA-insensitive *Rht-B1c* mutant, the RHT-B1C protein -which contains a 90 amino acid insertion in the N-terminal DELLA domain (16) -is not degraded after GA application (Figure 5C). In the *idd5* mutant, DELLA proteins are still degraded in response to GA, demonstrating that reduced stem elongation in this line is independent of RHT-1 degradation.

We next tested epistatic interactions between SLN1 and SDW3 by introducing the loss-of-function *sln1c* allele into selected *sdw3* backgrounds. In a ‘Himalaya’ background, the *sln1c* allele causes excessive elongation growth of all aerial parts and confers male sterility (18). When combined with the *sdw3a* allele, elongation growth was markedly reduced (Figure 5D, Table S12), and male fertility was restored, as indicated by a full grain set.

Although plant height was not significantly different between *sdw3a* and *sdw3a/sln1c* mutants, the *sdw3b/sln1c* mutant was significantly taller than *sdw3b* (Figure 5E, Table S12). This suggests that *sln1c* can partially rescue *sdw3b* to near wild-type height, although to a much smaller degree than in a wild-type background (18). Consistent with this, we observed slight elongation of the sub-crown internode in *sdw3b/sln1c* -a characteristic response of *sln1c* in a wild-type background -but not in the *sdw3a* background (Figure S9). These observations likely reflect that *sdw3b* (S40L) retains a low degree of SDW3 activity, compared to the more severe splice-site mutation in *sdw3a* (Table 1), consistent with a smaller reduction in height (see below).

### *idd5* and *sdw3* mutants exhibit a semi-dwarf phenotype

In glasshouse conditions, the *sdw3a* allele in barley reduced plant height by 24.3% compared to wild-type ‘Himalaya’ (Figures 6A and 6B, Table S13). The *sdw3e* allele conferred a similar reduction in plant height (22.6%), whereas *sdw3b* allele had a milder effect (10.3% reduction, Figure 6B, Table S13).

**Figure 6.**
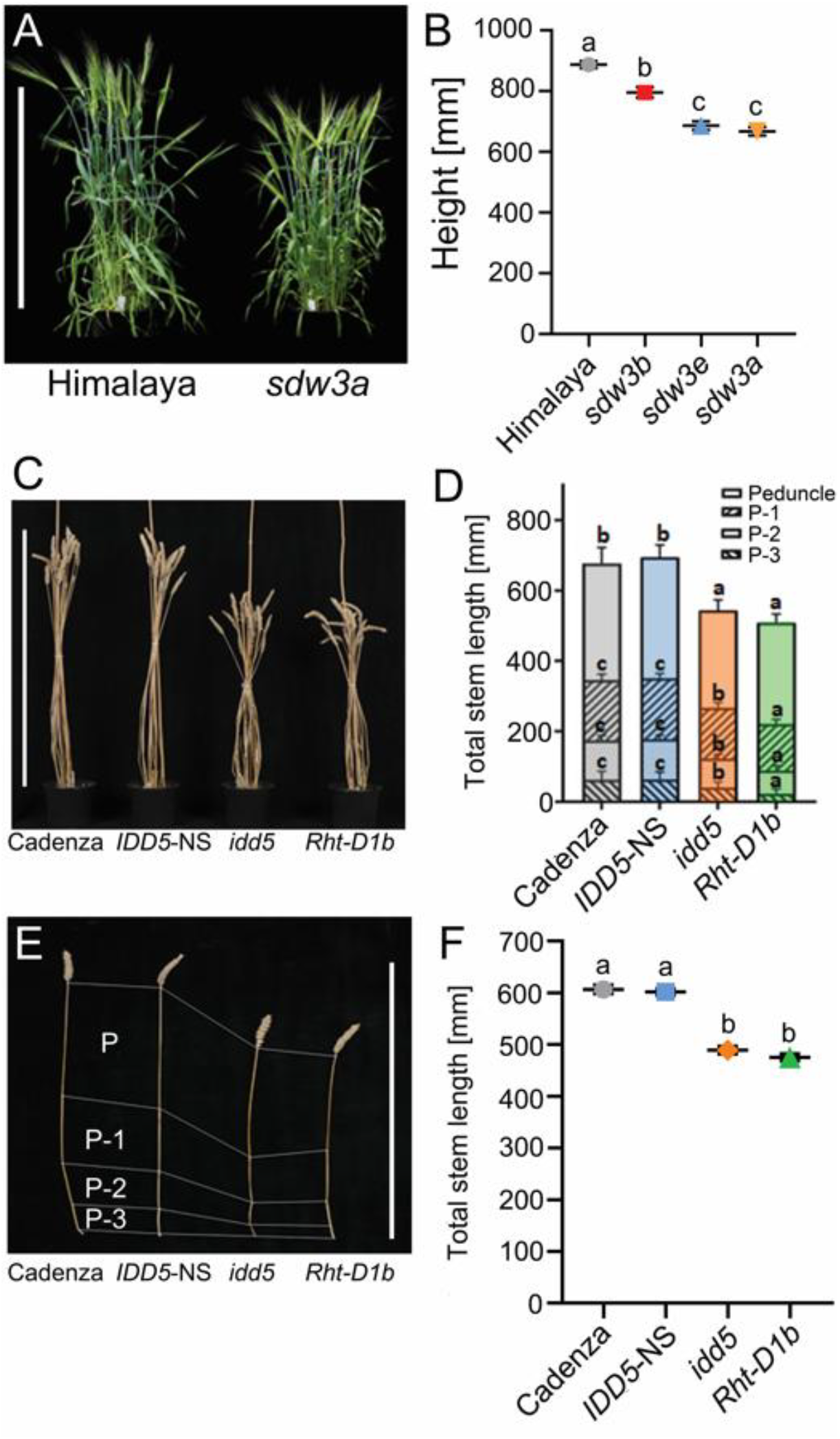
The phenotype of the mature barley *sdw3* mutants and wheat *idd5* mutants. **(A)** Mature phenotype of wild-type Himalaya and *sdw3a* (M671). **(B)** Mean height ± SEM of wild-type, *sdw3a, sdw3b* and *sdw3e* mutants (*N* = 11-12). Different letters indicate statistically significant differences determined by one-way ANOVA followed by Tukey’s post-hoc test (*P* < 0.05). **(C)** Mature phenotype of wheat genotypes grown in the glasshouse. **(D)** Mean length of representative tilers from each genotype with internodes indicated. P = peduncle, P-1 internode etc. **(E)** Contribution of each internode to the final stem length. **(F)** Analysis of the stem length data collected in the field trials in 2021. The wheat data, collected both from the glasshouse and field experiments, were analysed using one-way ANOVA and by linear mixed models followed by Tukey’s post-hoc LSD test (*P* < 0.05). Plotted values are means ± SEM. Bars in panels A, C and E are 80 cm.

The *idd5* mutant in wheat reduced total stem length by 21.5% compared to IDD5-NS (*P* <0.001) due to decreased elongation of all internodes (Figures 6C and 6D, Table S14). The effect was milder than the *Rht-D1b* allele which conferred a 25.9% reduction in stem length compared to ‘Cadenza’ (*P* < 0.001). All internodes in *idd5* except the peduncle were significantly longer than in *Rht-D1b* (Figure 6D, Table S14). Neither *idd5* nor *Rht-D1b* mutants developed a fifth internode, which was present in wild-type plants.

Field-grown *idd5* plants were 18.7% shorter than *IDD5*-NS, similar to the 21.6% reduction observed in *Rht-D1b* compared to ‘Cadenza’ (Figure 6F, Table S15). The height difference was mainly due to the reduced length of the uppermost internodes (Figure 6F, Table S15). Spike length was unaffected, but the *idd5* mutant produced fewer spikelets than *IDD5*-NS (Table S15).

## Discussion

Despite the importance of ‘Green Revolution’ DELLA alleles in modern agriculture, the transcriptional targets and precise mechanisms by which these proteins regulate growth and development remain poorly understood. Our discovery of IDD5 and SDW3 provides opportunities to characterise the mechanisms of DELLA-mediated control of TF activity, to identify the downstream regulatory networks controlling cell elongation in grasses, and to evaluate the potential benefits of alternative semi-dwarfing alleles for crop breeding.

### DELLA-mediated regulation of IDD activity

DELLA proteins regulate transcription networks indirectly through evolutionarily conserved interactions with multiple TF families (2). In Arabidopsis, at least seven members of the IDD family (including ENY and GAF1, homologues of IDD5 and SDW3, Figure S2) bind DELLA proteins (22). Furthermore, the interaction between AtIDD4 and DELLA proteins is conserved even in lycophyte and bryophyte lineages (2), suggesting that DELLA-mediated regulation of IDD proteins is an ancestral mechanism.

It will be important to determine how DELLA binding affects IDD transcriptional activity. One possibility is that DELLA proteins sequester or inhibit IDD5/SDW3, similar to their regulation of the positive growth regulator PIF4 (6, 7). In this model, the GA-induced degradation of DELLA proteins would relieve the inhibition of IDD5/SDW3, promoting growth. Alternatively, DELLA proteins may enhance IDD transcriptional activity by recruiting coactivator complexes (11), potentially to boost the expression of a growth inhibitor.

IDD proteins in the GAF1/ENY subclade, including SDW3 and IDD5, contain an EAR motif that mediates interaction with TPL repressors. The subsequent recruitment of chromatin-remodelling factors suppresses transcription of target genes (31). This regulation contributes to jasmonic acid (32) and auxin signalling (33, 34) and might also modulate GA signalling responses via IDD proteins. Consistent with this model, recent studies suggests that DELLA and TPL proteins can act as coactivators and corepressors of GAF1 and ENY (27, 28), although the regulatory mechanism was not fully elucidated. Distinguishing between these models will require identifying IDD5/SDW3 target loci and determining how DELLA activity modulates their transcription.

### IDD proteins regulate downstream pathways controlling cell elongation

Functional diversification among the wide range of DELLA-interacting TFs and their transcriptional targets likely drives the diversity in GA-regulated developmental responses. Understanding the different targets of IDD5, SDW3 and other DELLA-interacting proteins will enable more precise manipulation of DELLA-mediated growth and development. For example, PLATZ1, a positive regulator of cell elongation, also interacts with RHT1 (35). Understanding the degree to which PLATZ1 and IDD5 transcriptional targets overlap can help build our understanding of cell elongation in grass species.

IDD5 transcriptional targets might include GA biosynthesis genes. In *Arabidopsis*, the GAF1-GAI protein complex binds to the promoters of *AtGA20OX2, AtGA3OX1* and *AtGID1b* to induce their expression (25, 28). In peach (*Prunus persica* L. (Batsch)), an IDD1-DELLA complex activates *GA20OX1* expression (36) suggesting that IDD-DELLA complexes have a conserved role in GA homeostasis. In wheat, *GA3OX2* and *GA20OX* transcript levels were increased in *idd5* mutants, suggesting that this mechanism may also be conserved in cereals (Figure 4C).

The increased transcript levels of GA feedback-regulated genes and higher bioactive GA levels in the *idd5* mutant is unexpected given that IDD5 functions as a positive regulator of GA-responsive growth (Figure 4). One explanation is that DELLA proteins, released from sequestration in the absence of IDD5, may act as co-activators of GA biosynthesis genes with other transcriptional components. Other factors, such as the non-DELLA GRAS protein SCL3 (27), might also regulate these IDD targets. Given the opposing role of OsIDD2 as a negative regulator of GA-responsive cell elongation in rice (37), it is tempting to speculate that other IDD proteins may act in competition to regulate the transcription of these target genes. In barley, the IDD protein BROAD LEAF1 represses longitudinal cell division in leaves (38), a function shared by its Arabidopsis homologues AtIDD14, AtIDD15, and AtIDD16 (38). Similarly, both wheat and barley genomes contain paralogues of IDD5 and SDW3 that encode similar proteins containing an ID domain and EAR motif (Figure S2 and S3). In Arabidopsis, GAF1 and ENY paralogues exhibit partially redundant functions (24, 28), suggesting that IDD5 and IDD1 may similarly overlap in function in cereals. Generating additional loss-of-function mutants for these genes in wheat and barley will help test these hypotheses.

It will also be important to study the effects of IDD5 and SDW3 activity in different tissues. In *Arabidopsis*, GAF1 and ENY regulate seed germination and starch accumulation (24), so examining these traits in wheat and barley could reveal additional phenotypes affected by these genes. Although *idd5* mutations restore the height phenotypes of the *rht-1* loss-of-function mutants, they do not rescue fertility, indicating that other RHT1-interacting proteins are likely responsible for GA signalling in male fertility. It is interesting to note that *sdw3* mutations overcome sterility in barley *sln1c* loss-of-function mutants, suggesting either the functional divergence of SDW3 and IDD5 or the involvement of different IDD proteins in reproductive development in each species.

### Application of *IDD5* and *SDW3* alleles in agriculture

In both glasshouse and field trials, *idd5* and *sdw3* loss-of-function mutants reduced plant height to a degree comparable to the ‘Green Revolution’ *Rht-D1b* allele. Moreover, the ability of *idd5* and *sdw3* mutations to fully rescue the excessive elongation of *della*-null mutants demonstrates the central role of IDD5/SDW3 TFs in regulating stem elongation in barley and wheat. Because DELLA proteins remain responsive to GA-mediated degradation in these lines (Figure 5), these alleles could provide the benefits of semi-dwarf stature without the negative pleiotropic effects often seen with the ‘Green Revolution’ dwarfing alleles, such as smaller grain size, poor seedling emergence, and lower nitrogen use efficiency (Figure S10). Comprehensive yield-plot phenotyping across multiple genotypes and environments will be necessary to evaluate the agronomic performance of these alleles.

Analysis of 44 sequenced barley pangenome accessions shows they all carry *SDW3* alleles encoding full-length proteins with intact functional domains (Table S16) (39). This suggests that *sdw3* loss-of-function mutations have not been widely exploited in barley breeding. Introducing these alleles using CRISPR/Cas9-based gene editing could, therefore, be a straightforward approach to engineer novel semi-dwarfing traits. Additionally, screening diverse germplasm collections might uncover additional natural *SDW3* alleles, including those -such as *sdw3b* -that retain some function and confer milder height reductions (Figure 6B). Such variation could allow breeders to tailor height reductions to specific environments or agronomic requirements.

It is unlikely that *idd5* alleles are present in modern wheat breeding lines due to the functional redundancy among the three homoeologous copies. Achieving a beneficial semi-dwarf growth habit likely requires mutations in all three *IDD5* homoeologous, since plants carrying loss-of-function mutations in one or two homoeologues exhibit only mild reductions in height (Figure S11, Table S17). However, selecting for three loss-of-function alleles will be more complex for breeders than deploying a single dominant gain-of-function mutation such as *Rht-D1b*.

## Materials and Methods

### Plant materials and growth conditions

All wheat lines were in the ‘Cadenza’ background. Near-isogenic lines carrying *Rht-D1a* and *Rht-D1b* alleles were described previously (17). EMS-mutagenized M_4_ lines carrying *IDD5* or *RHT1* mutations were identified from an *in silico* database (40). M_4_ plants of each *IDD5* mutant line were inter-crossed and then backcrossed three times to ‘Cadenza’. From the resulting BC_3_F_2_ populations, *IDD5*-NS and *idd5* genotypes were selected using assays described in Table S18. To determine the effect of the intron 1 donor splice site mutation in *IDD5-B1*, PCR amplicons spanning exons 1 and 2 of all three *IDD5* homoeologues were amplified using primers described in Table S18. Illumina sequencing libraries were generated from these pooled amplicons and sequenced at the Rothamsted Research sequencing core. Reads were aligned to *IDD5* gene models using Bowtie 2, and the proportion of correctly versus incorrectly spliced reads were calculated from read pileups (Figure S5B).

Similarly, M_4_ plants from each *RHT1* mutant line were backcrossed twice to ‘Cadenza’ and then crossed to combine the mutations, producing BC_2_F_2_ lines homozygous for knockouts in *RHT-A1* and *RHT-B1* and heterozygous for the *RHT-D1* mutation. These lines were maintained in a heterozygous state in *RHT-D1* to overcome the sterility of the *rht1*-null mutant. BC2F3 plants homozygous for all three *RHT1* knockouts were used in subsequent experiments. Crosses between *rht-1* and *idd5* mutants generated the sextuple *idd5/rht-1* line.

Initial phenotyping was performed in the glasshouse at 20°C using a 16 h/8 h light/dark cycle. All other experiments were conducted in controlled environment (CE) growth conditions under a 16 h photoperiod (300 µmolm_-2_s_-1_) with day/night temperatures of 20 °C/16 °C.

All barley materials were in the ‘Himalaya’ background. To identify *SDW3* mutants, wild-type ‘Himalaya’ grains were mutagenised with sodium azide (41) and then sown in the field. Bulk M_2_ grains were harvested and screened in soil in wooden flats that were watered with 1 µM GA_3_ solution. Most seedlings were highly elongated and pale-green in response to GA_3_ treatment but approximately 1 in 1,000 seedlings were dark-green and semi-dwarf. These semi-dwarf plants were transplanted to pots and grown to maturity without further GA treatment. Inter-crossing of these lines over subsequent generations indicated that they included several different genetic loci. Ten mutants defined by M671 formed a single complementation group (30). Further complementation tests with previously described dwarf barley mutants showed that M671 is allelic to the dwarfing mutation in Hv287, originally designated *GA-insensitive* (*GA-ins*) (42) but now renamed *Sdw3* (43). All *sdw3* alleles were confirmed by Sanger sequencing of three overlapping PCR amplicons (Table S18). In the Himalaya x M671 and *sdw3a* x *sln1c* populations, *sdw3a* was genotyped by Sanger sequencing the amplicon generated from primers Sdw3_F1/Sdw3_R1 (Table S18) and *sln1c* was genotyped by Sanger sequencing of PCR amplicons as described previously (18). Other barley mutants used in this study included *gse1a* (M488), which carries an amino acid substitution in the HvGID1 GA receptor (44, 45), *grd2b* (M463), which is GA-deficient due to a loss-of-function mutation in *HvGA3OX2* and *sln1c* (18). All barley plants were grown under standard glasshouse conditions at 20°C with a 16 h/8 h light/dark cycle.

### Field experiment

Wheat lines were sown in spring 2021 at the Rothamsted Experimental farm in Southeast England in 1 m_2_ plots arranged in a randomised block design at a rate of 450 seeds/m_2_. The field trial plan comprised 108 1 m_2_ plots arranged in three replicate blocks of 6 rows and 6 columns, with the complete trial arranged as 18 rows by 6 columns. Lines (39 in total) were allocated to plots according to a resolvable row-column design with blocking as described and additional Latinization across ‘long columns’ (i.e. single columns of 18 plots cutting across the replicate blocks) to allow for potential additional variation due to the direction of farming operations (perpendicular to spatial blocking). Standard farm practice for fertiliser and pesticides was followed, with no plant growth regulators applied. Height measurements were made using a floating polystyrene disc, taking six measurements per plot. Internode lengths were measured for ten individual tillers by measuring between the lower bounds of each node. Grain sizes were measured on samples of ∼200 grains using a MARVIN-Digital Seed Analyser (MARViTECH GmbH, Germany). Data were analysed using linear mixed models fitted using restricted maximum likelihood, each with a one-way fixed model (line) and the random model reflecting the design/randomization structure of the trial ((block/row)×long column). Whilst data for all lines grown were included in the analysis results for comparisons between only the four lines of interest are given here (extracted using a nested treatment contrast).

### Cell length measurements

Two images of abaxial epidermis cells were taken in the uppermost 10 mm of the fully elongated leaf sheath using a JEOL JSM-6360LV scanning electron microscope. Twelve plants per genotype and treatment were analysed. All visible individual cells in each image were measured with ImageJ 1.48v software (46). The data were compared using unbalanced ANOVA with Tukey’s post-hoc HSD test.

### GA dose-response assays

Wheat seeds were surface sterilised, germinated and transplanted into vermiculite saturated with water or GA_3_ solutions ranging from 1 nM to 100 µM in ten-fold increments. Plants were treated every two days for ten days, after which seedlings were harvested and the first leaf sheath and blade lengths were measured. Treatments were replicated three times in a blocked design. Barley seeds were germinated in filter paper envelopes soaked with either water or GA_3_ solutions. The maximum leaf elongation rate (LER_max_) of each seedling was determined following the method described previously (45). Genotypes with a normal GA response reach near-saturation of leaf elongation at GA_3_ concentrations of 1 – 10 µM (45). By contrast, the *gse1* mutant is less sensitive to GA and shows continued increases in leaf elongation at very high (mM) concentrations of GA_3_ (44). All measurements were analyzed using one-way ANOVA, accounting for replication and blocking in a nested treatment structure (Block/Unit). The least significant difference (LSD) was set at the 5% level. GenStat (20th edition, 2019, ©VSN International, Hemel Hempstead, UK) was used for all analyses, and residual and mean plots were examined to confirm data normality.

### Protein-protein interaction assays

A wheat cDNA library was constructed from RNA extracted from mature aleurone tissues. Grains were de-embryonated and incubated for 3 d in 20 mM CaCl_2_, after which the aleurone layers were isolated from half-grains. Total RNA was extracted and used to generate cDNA prey libraries (Life Technologies Corporation). This library was screened using a plasmid containing an N-terminally truncated RHT-D1 fragment (Δ*RHT-D1*, nucleotides 652 -1872 bp, amino acids 218-623) described previously (16). To transform the library, 250 μl of library-scale MaV203 competent cells (>1×10_6_ transformants, Thermo-fisher Scientific, California, USA) was mixed with 10 μg of bait plasmid, 10 μg of wheat aleurone cDNA prey library and 1.5 ml of PEG/LiAc solution. The mixture was incubated at 30 °C for 30 min, then treated with 88 μl DMSO (Sigma-Aldrich, Darmstadt, Germany) and heat-shocked at 42°C for 20 min. Pairwise interactions were tested in a Y2H system with plasmids expressing *IDD5-A1*, Δ*SLN1* (640 – 1857 bp), *SDW3*, and truncated forms of *TaIDD5-A1* (M1, M2, M3 and M4). Plasmids were introduced into *MaV203* yeast competent cells by heat shock. 150 µl of competent yeast cells were incubated with 1 µg of each plasmid DNA, 2 µl of 10 mg/ml sheared salmon sperm DNA (Thermo-fisher Scientific, California, USA) and 350 µl of 50% polyethylene glycol (PEG3350) at 30 °C for 30 min, then 42 °C for 5 min, placed on ice for 2 min and centrifuged. Cell pellets were resuspended in 110 µl of sterile distilled water and plated on SD-Leu-Trp selective media, then incubated at 30 °C for 72 h. A small amount of yeast colony originating from a single cell was resuspended in 200 µl of sterile water. A 5 µl aliquot of this suspension was spotted onto SD-Leu-Trp-His medium containing 50 and 100 mM of 3-AT and onto SD-Leu-Trp-Ura medium. Three biological replicates were performed for each strain and plates were incubated at 30 °C for 72 h to assess relative growth.

For bimolecular fluorescence complementation, destination vectors (AB830561, AB830564, AB830568, AB830572) used in this study were described previously (47). Full-length *RHT-D1* and *IDD5-A1* coding sequences were inserted into these vectors using Gateway cloning. The *IDD5-A1* gene was synthesised by GenScript (Netherlands), and the codons optimised for expression in *Nicotiana benthamiana*. Cultures of *Agrobacterium tumefaciens* strain GV3101 carrying the fusion constructs were grown overnight in 2YT media containing 50 µg/ml rifampicin, 25 µg/ml gentamicin and 100 µg/ml spectinomycin. One ml of each culture was pelleted (1,505 x g for 5 min), resuspended in infiltration medium (28 mM D-glucose, 50 mM MES, 2 mM Na_3_PO_4_·12H_2_O, 100 µM acetosyringone) to an OD_600_ of 0.1, and mixed 1:1:1 with a p19 control culture. The resulting suspensions were infiltrated into the abaxial sides of leaves from six- or seven-week-old *N. benthamiana* plants. After 3 d, infiltrated leaves were examined using a Zeiss LSM 780 confocal microscope with Leica Application Suite X (LAS X) software.

### Western blot

Proteins were extracted from developing wheat spikes by homogenizing the tissue in an extraction buffer containing one cOmplete™ ULTRA Mini EDTA-free Protease Inhibitor Cocktail Tablet (Merck KGaA, Darmstadt, Germany), 2 ml of 5x lysis buffer (50 mM Tris/Cl pH 7.5, 750 mM NaCl, 2.5 mM EDTA) and 100 µl of 100 mM PMSF. After a 20 min incubation, samples were denatured at 70 °C for 10 min, separated by SDS-PAGE, and transferred onto 0.2 µm PVDF membranes. Membranes were first stained with Ponceau S to confirm equal protein loading, then blocked and incubated overnight (16 h at 4 °C) with rabbit anti-SLR1 primary antibody (1:5000). Following washes, membranes were incubated for 4 h at 4 °C with goat anti-rabbit IgG alkaline phosphatase (1:5000 dilution, Merck) before developing with the BCIP®/NBT-Purple Liquid Substrate System (Merck) for up to 2 h.

### GA hormone extraction and analysis

Gibberellin levels were quantified in the leaf sheath tissue of ‘Cadenza’, *IDD5*-NS, *idd5* and *Rht-D1b* genotypes, using four biological replicates per genotype. Seven-day-old seedlings were harvested from the grain to the top of the coleoptile, and the leaf sheaths were freeze-dried. GAs were extracted from ground tissue homogenate and purified with an Oasis® MAX, mixed mode SPE column (Waters, Ireland). An Aquity UPLC I-Class system (Waters, USA) was used to separate different GAs, which were directly detected using multiple-reaction monitoring mode (MRM) by triple stage quadrupole mass spectrometer Xevo TQ-XS (Waters, USA) equipped with electrospray interface working in negative mode (48) Masslynx 4.2 software (Waters, USA) was used to analyze the data and the quantitation of GA levels was performed based on the standard isotope dilution method (49). Data was the statistically processed using one-way ANOVA and Tukey’s post-hoc HSD test.

### Transcriptome analysis

Leaf sheaths from seven-day-old ‘Cadenza’, *idd5* and *Rht-D1b* plants (four biological replicates each) were collected 8 h after treatment with either water or 100 µM GA_3_. Tissue from the grain to the top of the coleoptile was harvested, flash-frozen in liquid nitrogen and total RNA was extracted using the Monarch® Total RNA Miniprep Kit with on-column DNase treatment (New England Biolabs, Ipswich, Massachusetts, USA). RNA quality was assessed using an Agilent RNA 6000 Nano Chip and Agilent 2100 Bioanalyzer (Agilent, Santa Clara, USA).

RNA samples were sequenced using paired-end Illumina sequencing by Novogene Europe (Cambridge, UK). Sequencing reads were trimmed for quality and adapter sequences using Trimmomatic 0.39 (parameters SLIDINGWINDOW:4:20; MINLEN:50) (50), then aligned and quantified using Kallisto against the IWGSC RefSeq v1.2 annotated gene models (51, 52). Transcript per million (TPM) values were calculated for each gene and pairwise contrasts were made between control and GA_3_ treatments for each genotype, as well as between genotypes for the same treatment using DESeq v2 (53). UpSet plots (54) were used to visualize DEGs, and TPM values of selected genes were displayed with the online tool “Heatmapper” (http://www.heatmapper.ca/expression/).

Total RNA was extracted from young barley shoots grown in filter paper envelopes and sequenced as 2 x 150 bp paired end Illumina sequencing by Novogene (China). Trimmomatic (50) was used to remove low-quality bases and adapter sequences, and the resulting high-quality paired reads were mapped to Morex v2 barley reference gene models (55) using Tophat2/Bowtie2 tools (bowtie/2.3.4 and tophat/2.1.1; (54) with default parameters. Sorting BAM files and removing duplicate reads was performed as described previously (56). Single nucleotide polymorphism (SNP) calling and candidate gene identification were performed using the MUTRIGO pipeline (https://github.com/TC-He witt/MuTrigo) with default settings.

### Phylogenetic analysis

Putative IDD family members in wheat and barley were identified using BLASTp searches against the IWGSC RefSeq v1.2 and Morex V3 proteome databases through the Ensembl Plants platform (https://plants.ensembl.org/index.html). Rice IDD family members and the INDETERMINATE domain were used as search queries. Phylogenetic analysis was conducted in PhMyl (57) with a BLOSUM62 substitution model and 100 bootstrap replicates. Protein alignments were produced using MUSCLE (58). SDW3 proteins were identified using BLASTp from the proteomes of 44 barley accessions from the barley pangenome (39).

## Acknowledgements

This paper is dedicated to our friend and colleague, Matthias, who passed away before this work was published. This project was funded by the Biotechnology and Biological Sciences Research Council (BBSRC) through the Delivering Sustainable Wheat (BB/X011003/1) and Designing Future Wheat (BB/P016855/1) programmes. PH and DT acknowledge funding from the European Regional Developmental Fund Project ‘Towards Next Generation Crops’ No. CZ.02.01.01/00/22_008/0004581, www.tangenc.cz).

