## Supplementary_figures_and_tables for "INDETERMINATE DOMAIN-DELLA protein interactions orchestrate gibberellin-mediated cell elongation in wheat and barley"

**Figures S1 to S11**
**Tables S1 to S18**

**Figure S1.** RNA-seq read coverage of wild-type 'Himalaya' and eight *sdw3* mutants across the genomic sequence of *HORVU.MOREX.r3.2HG0131300* (*SDW3*). Polymorphisms with the reference sequence are highlighted with stars. M671 (*sdw3a*) has a SNP in the intron 1 splice donor site causing aberrant splicing and a shift in read coverage across the first intron which is highlighted with a blue bar. Sequences of all alleles are provided in additional file 1.

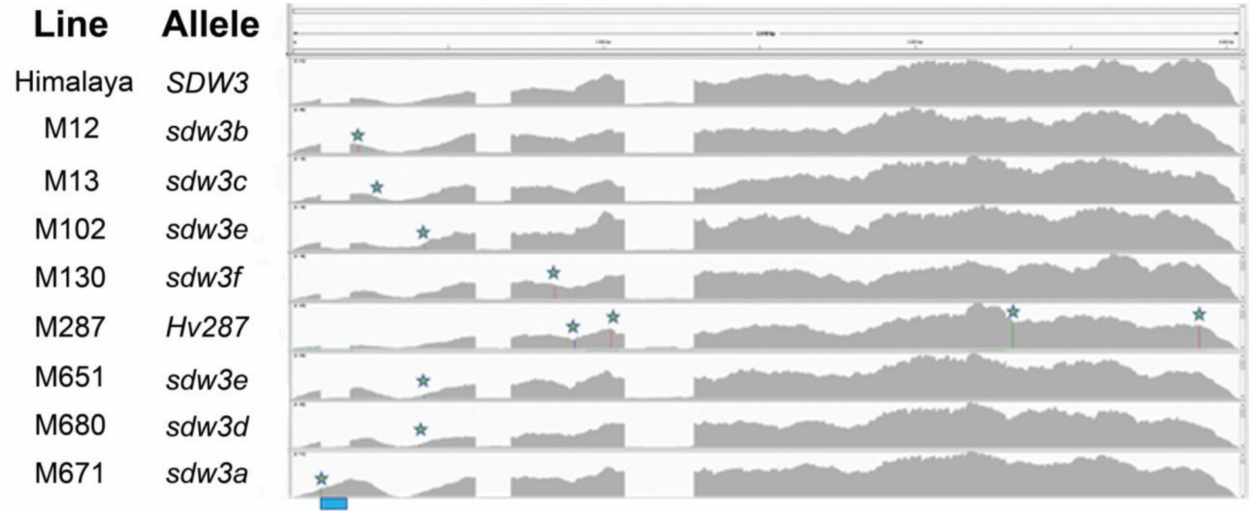

**Figure S2.** Phylogenetic tree of IDD proteins in Arabidopsis (red), rice (blue), barley (green), and wheat (black). Phylogenetic analysis was conducted using a BLOSUM62 substitution model with 100 bootstraps. The ENY/GAF1 subclade is indicated.

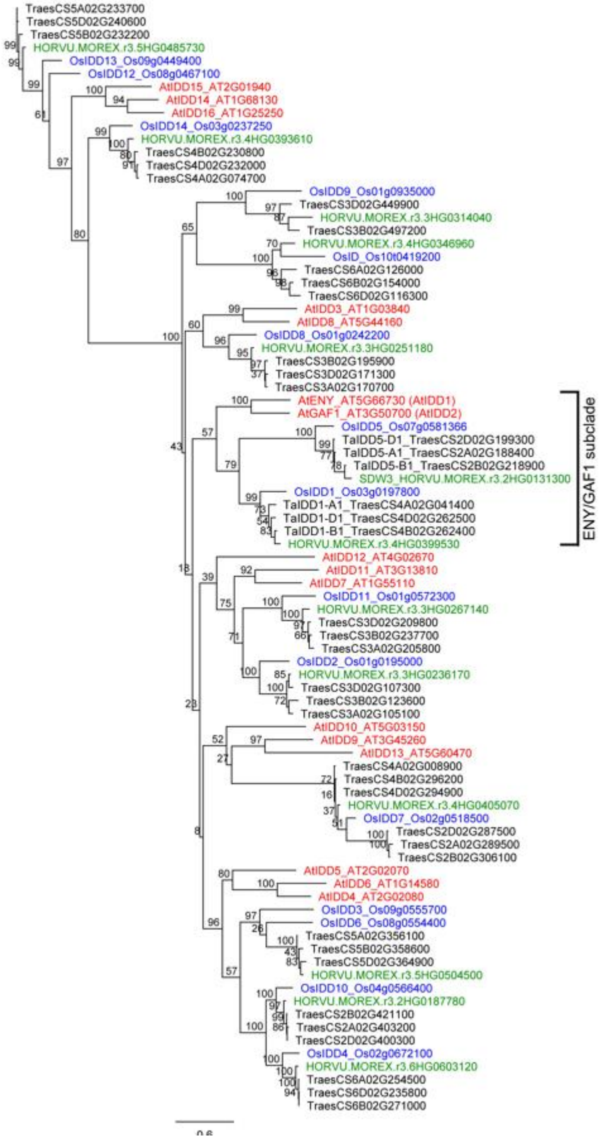

**Figure S3.** Amino acid alignment of Arabidopsis, wheat, barley and rice IDD proteins in the ENY/GAF1 subclade. The conserved ID domain, PAM domain and ERF motif are indicated. The position of *SDW3* alleles that encode non-synonymous amino acid substitutions and insertions are shown.

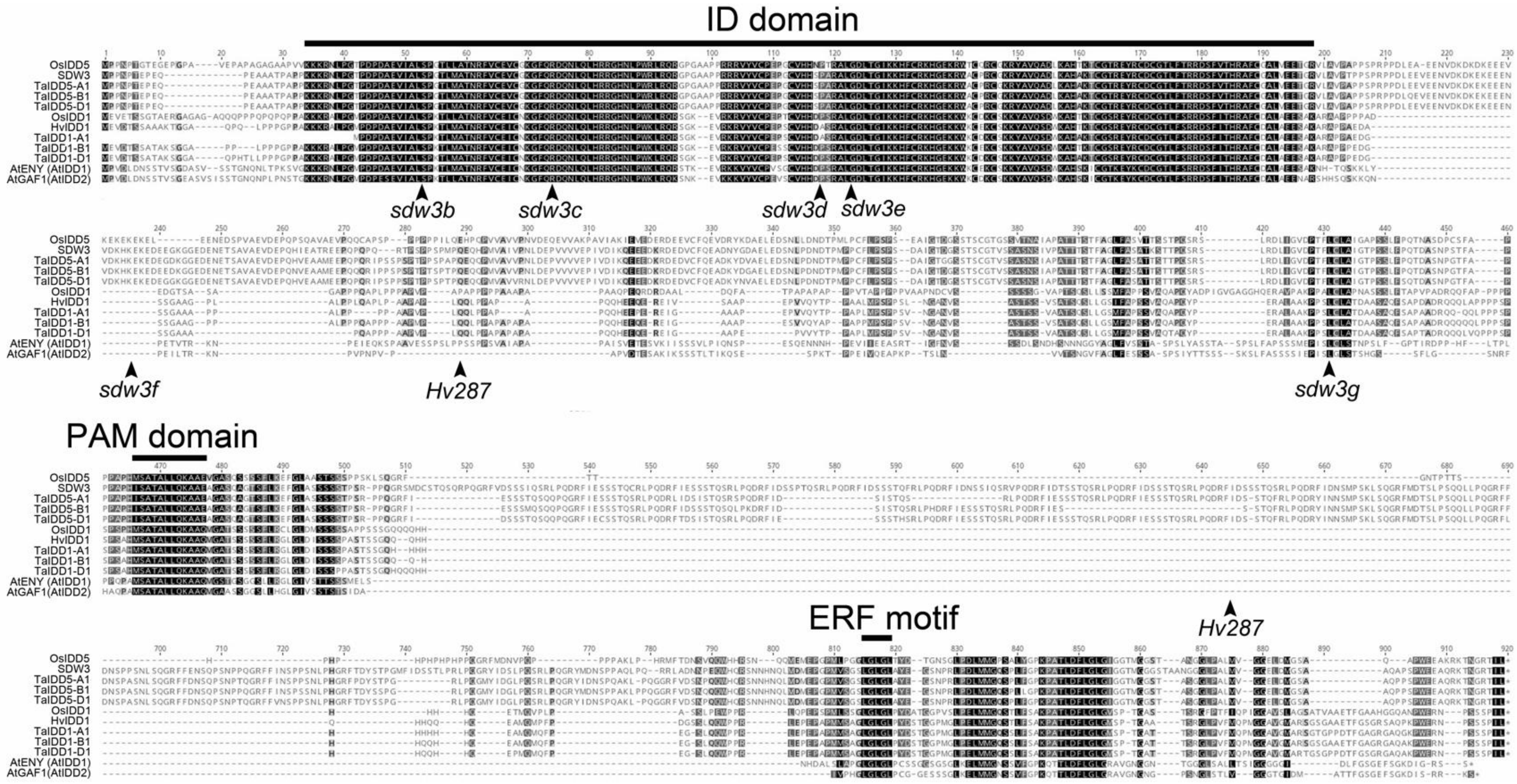

**Figure S4**. Pairwise amino acid identity (%) between full-length TaIDD5 and HvSDW3 protein sequences.

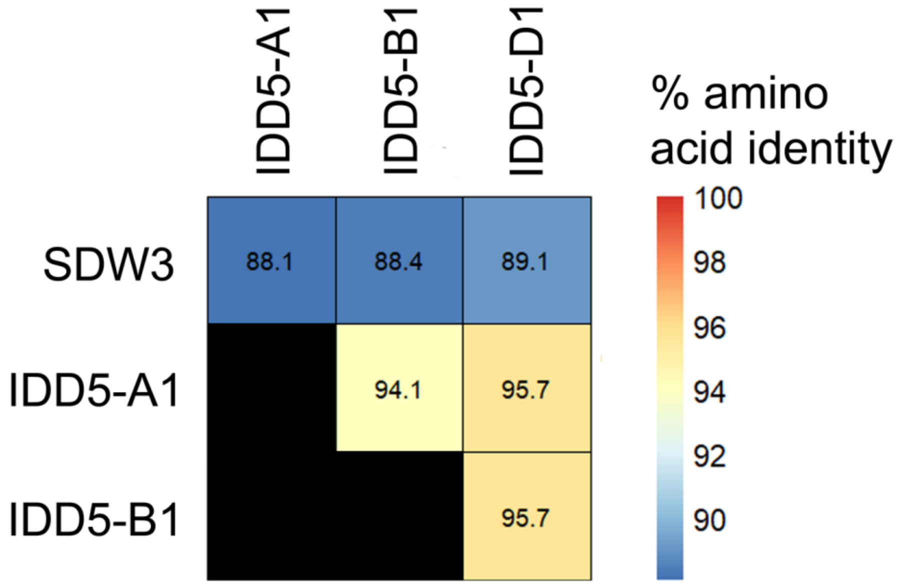

**Figure S5. (A)** Position of EMS-induced mutations in *IDD5-A1*, *IDD5-B1* and *IDD5-D1* used in this study. **(B)** Proportion of spliced and unspliced intron 1 reads in *IDD5-B1* in wild-type and CAD4-1415 lines. The proportion of reads mapping to each homoeologue was calculated manually from read pileups.

A

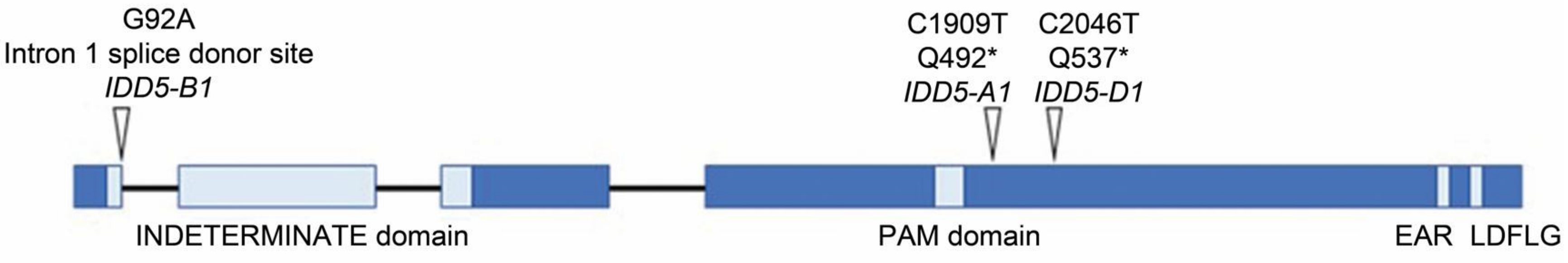


B

| Genotype | % spliced reads | % unspliced reads | Contribution of homoeologues to unspliced reads | |
|---|---|---|---|---|
| Wild-type | 92% | 8% | *IDD5-A1* | 28% |
| | | | *IDD5-B1* | 40% |
| | | | *IDD5-D1* | 32% |
| CAD4-1415 | 65% | 35% | *IDD5-A1* | 7% |
| | | | *IDD5-B1* | 86% |
| | | | *IDD5-D1* | 7% |

**Figure S6.** GA dose response curve in L1 leaf blade tissues of seedlings 14 days after germination. Data were analysed using one-way ANOVA and Tukey's post-hoc HSD test.

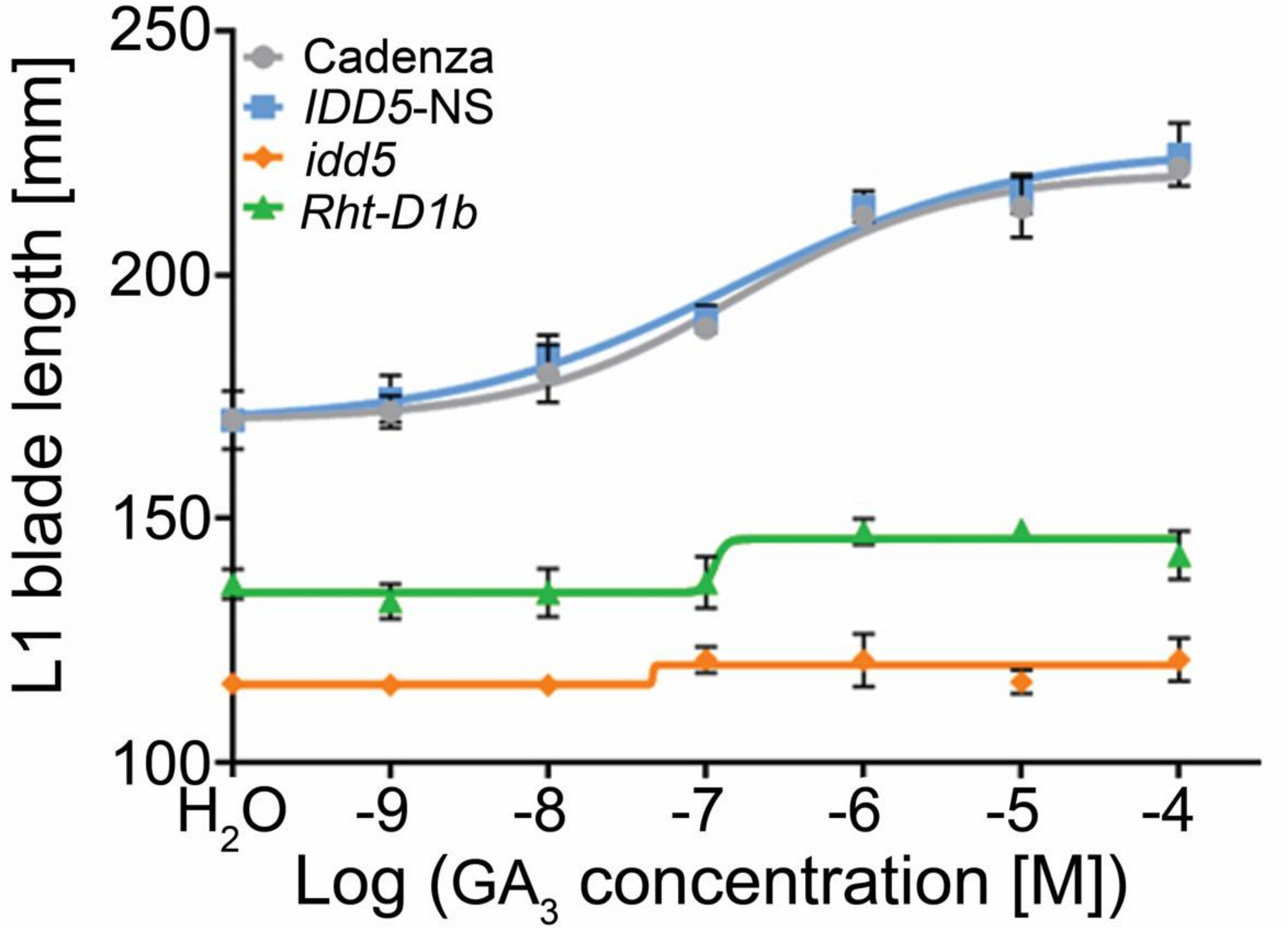

**Figure S7.** L1 sheath length (mm) of seven-day-old ‘Cadenza’ and *idd5* seedlings without (-GA) and after 100 µM $GA_3$ treatment (GA). Data were analysed with one-way ANOVA and Fishers post-hoc HSD test. Different letters indicate significant differences between genotypes at the 0.05 confidence level.

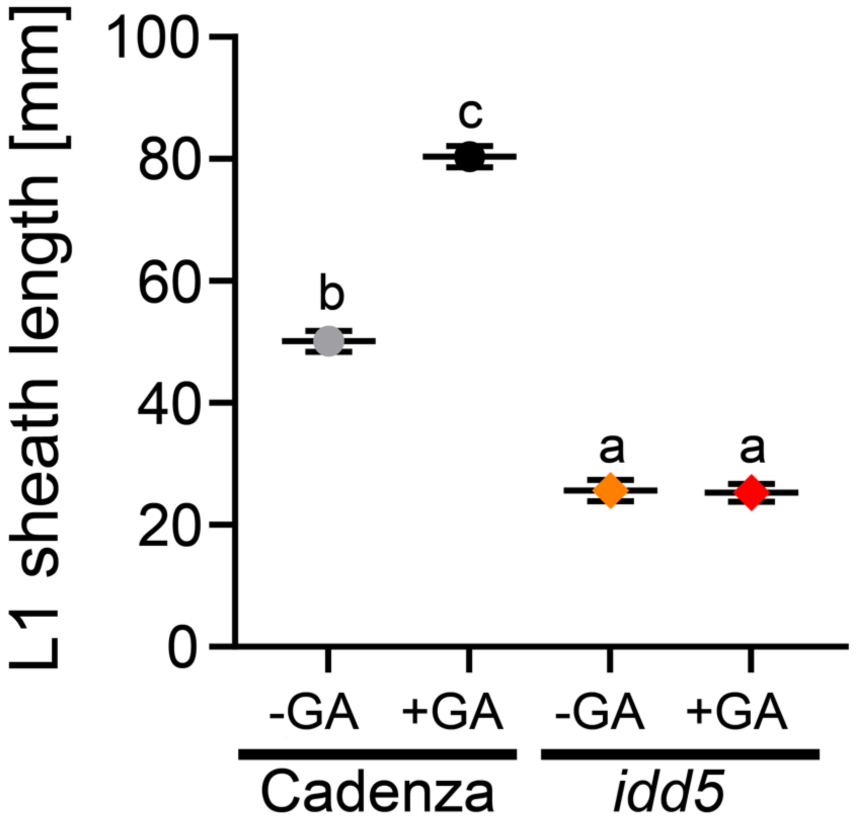

**Figure S8**. Principal Component Analysis plot of RNA-seq data generated from Cadenza, *Rht-D1b* and *idd5* lines.

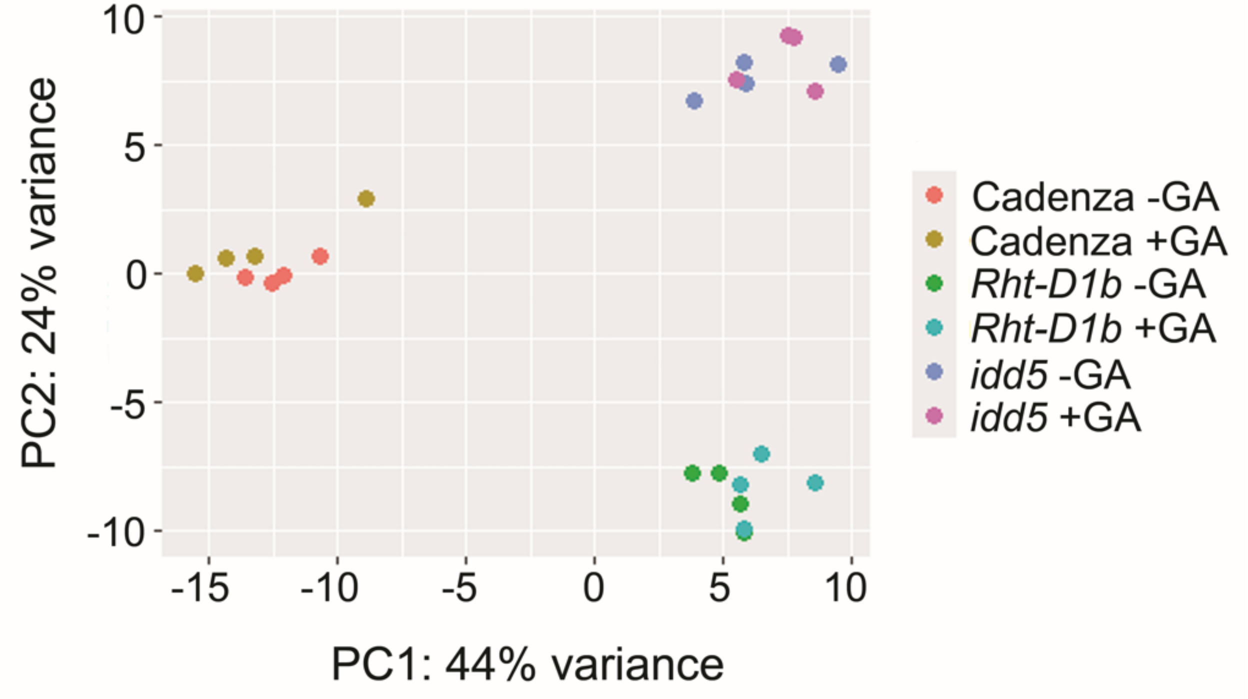

**Figure S8**. UpSet plot showing the number of DEGs in selected pairwise comparisons ($P_{adj}$ < 0.01) and the subset of those genes that are common to different contrasts in GA-treated tissues of Cadenza, *Rht-D1b* and *idd5* genotypes.

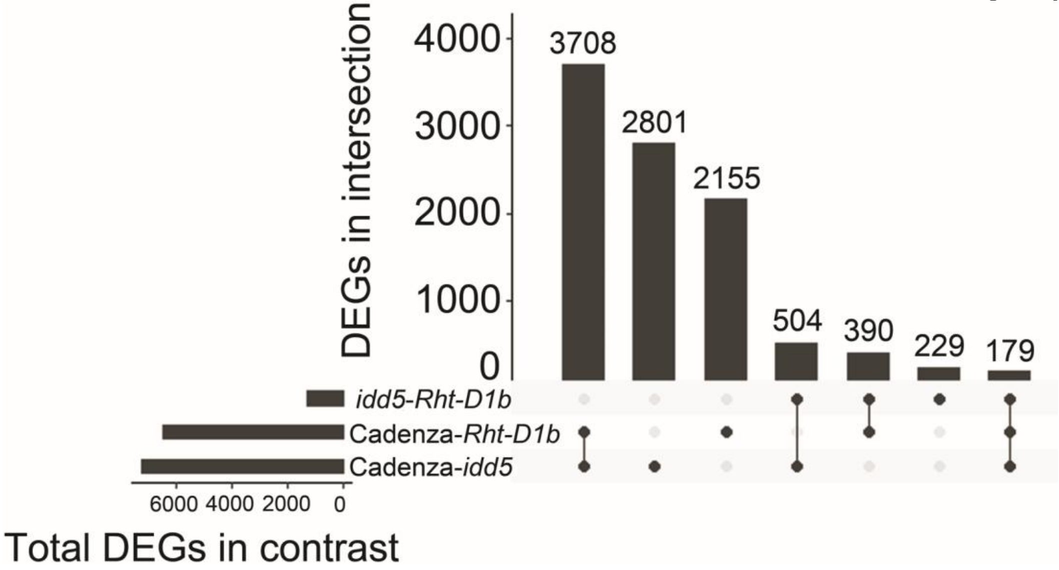

**Figure S9.** Representative images of the sub-crown internode (SCI) in *sdw3b* and *sdw3b/sln1c* double mutants.

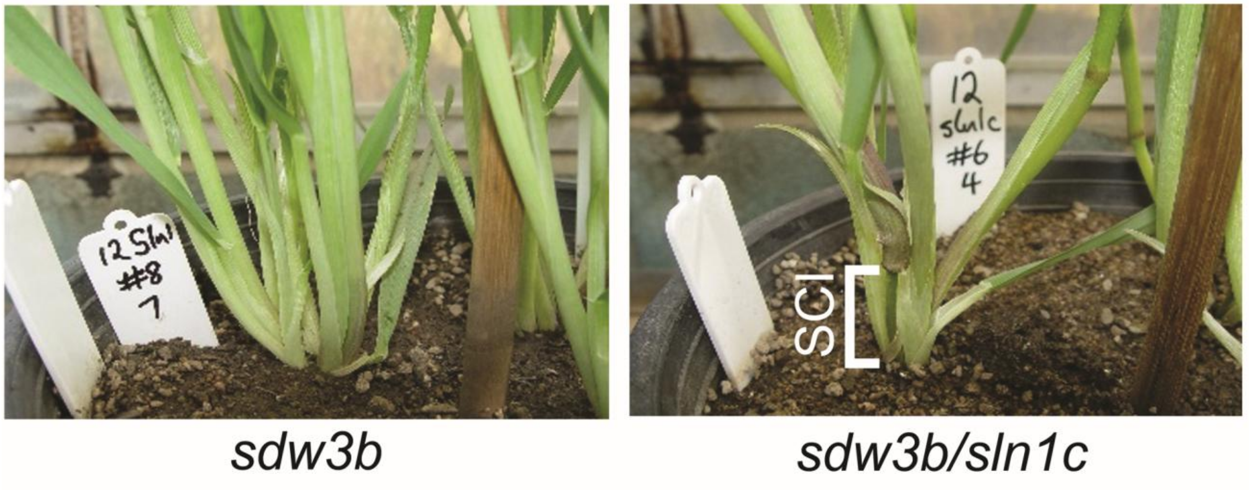

**Figure S10.** Potential benefits of alternative dwarfing alleles in crops. In wild-type plants, GA-mediated degradation of DELLA proteins modulates the activity of hundreds of downstream transcription factors, including IDD5 and SDW3, to coordinate growth and development. The *Rht-D1b* allele selected during the 'Green Revolution' encodes a GA-insensitive DELLA protein that constitutively affects the activity of all downstream transcription factors, resulting in a beneficial semi-dwarf phenotype in addition to some undesirable negative pleiotropic effects. Targeting specific transcription factors, such as IDD5/SDW3, through loss-of-function mutations can reduce stem elongation while minimizing the negative pleiotropic effects associated with gain-of-function DELLA alleles.

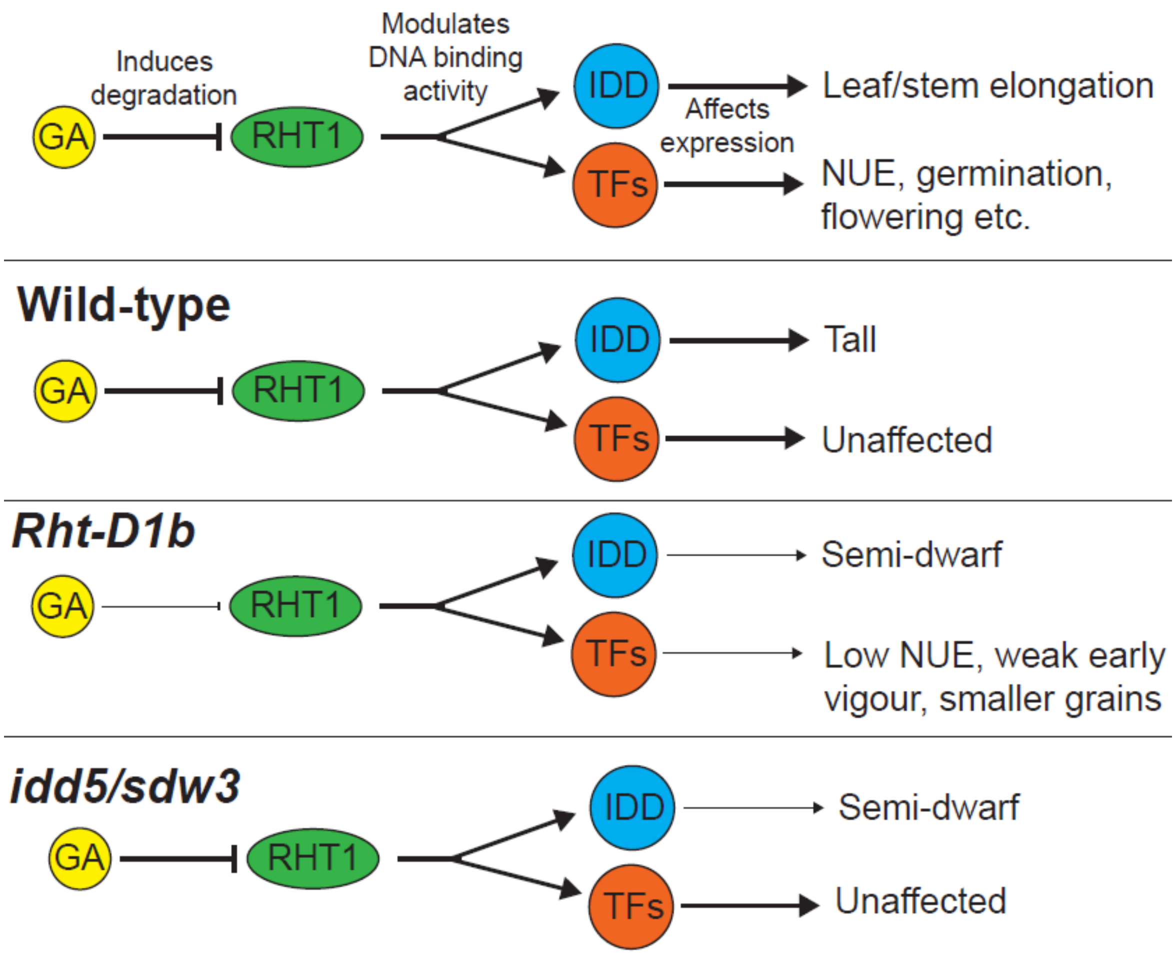

**Figure S11.** Stem length (mm) of single, double and triple *idd5* mutant genotypes in glasshouse conditions. Wild-type alleles are denoted in upper case and mutant alleles in lower case. Values are means ± SEM. Data were analysed with one-way ANOVA and Fishers post-hoc HSD test. Different letters indicate significant differences between genotypes at the 0.05 confidence level.

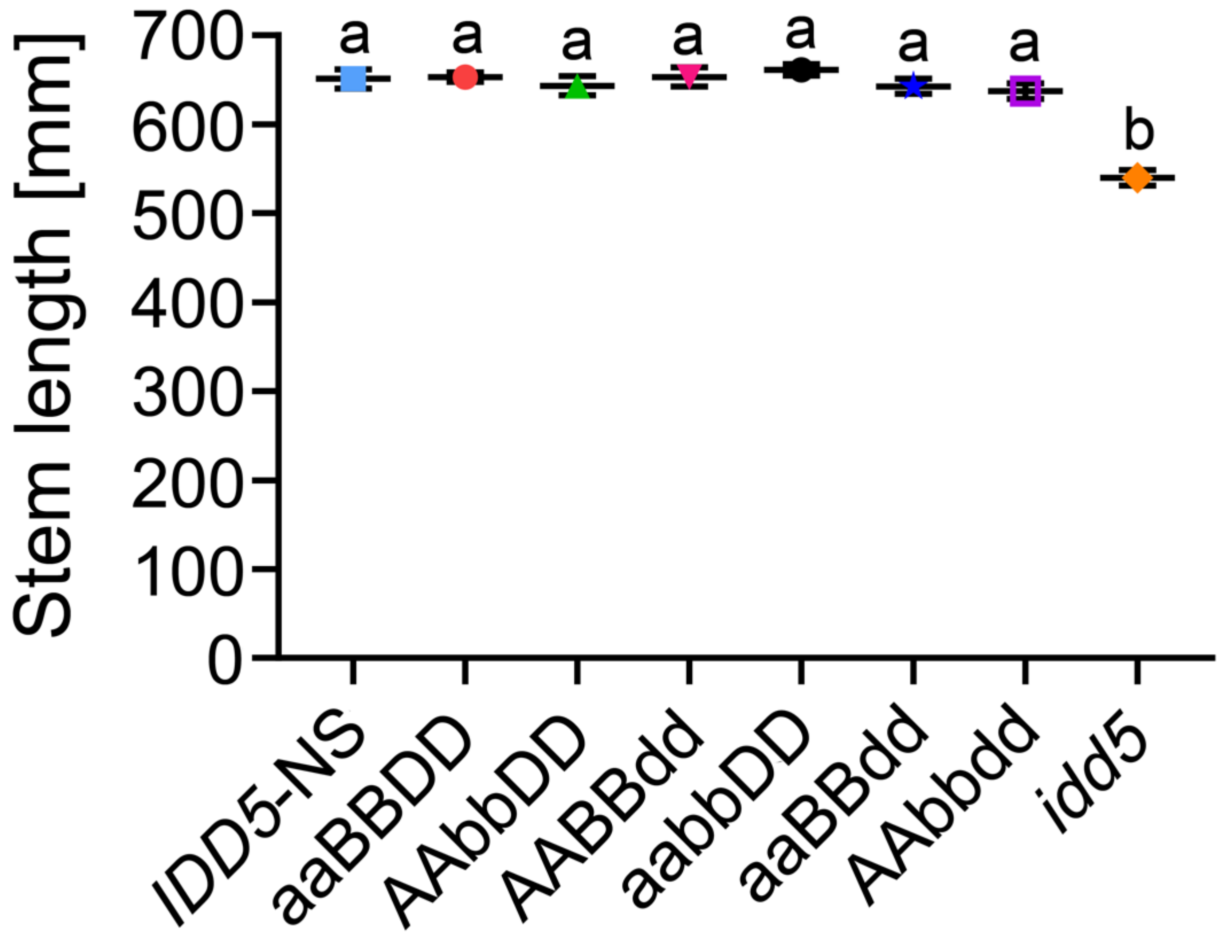

**Supplemental tables**

**Table S1.** Physical locations of *SDW3* markers on chromosome 2H of the Morex v2 and v3 barley genome assemblies. * = Previously identified gene-flanking markers described in Vu *et al*., 2010.

| | **Position on chromosome 2H (bp)** | |
|---|---|---|
| **Marker** | **Morex v2** | **Morex v3** |
| TC146454 | 133,393,757 | 132,621,149 |
| TC149567* | 134,398,749 | 133,603,150 |
| TC147542 | 136,535,222 | 135,675,412 |
| TC144440 | N/A | 137,048,490 |
| TC149841 | 139,015,090 | 138,038,964 |
| *SDW3* (*HORVU.MOREX.r3.2HG0131300*) | 140,488,530 | 139,466,760 |
| TC142185* | 140,491,098 | 139,469,633 |
| HW01K11(S2279) | 141,321,401 | 140,285,297 |

**Table S2**. Maximum leaf extension rates ($LER_{max}$) in a Himalaya x M671 $F_2$ population segregating for the *sdw3a* allele. Values are means ± SEM. Data was analysed using unbalanced two-way ANOVA. *P*-values represent the contrast between control and GA-treated seedlings from each genotype.

| **Genotype** | **$LER_{max}$ (mm $d^{-1}$)** | | ***P*-value** |
|---|---|---|---|
| | **Control** | **+ 10 μM $GA_3$** | |
| Himalaya (*SDW3*/*SDW3*) | 28.5 ± 1.2 (*N* = 10) | 51.7 ± 2.3 (*N* = 3) | < 0.001 |
| Heterozygote (*SDW3*/*sdw3a*) | 27.9 ± 0.7 (*N* = 28) | 36.2 ± 0.8 (*N* = 27) | < 0.001 |
| M671 (*sdw3a*/*sdw3a*) | 21.3 ± 1.2 (*N* = 10) | 21.7 ± 1.1 (*N* = 13) | 0.812 |

**Table S3**. Maximum leaf extension ($LER_{max}$) rates in Himalaya, *gse1a* and *sdw3a* genotypes in response to treatment with low (10 μM) and high (10 mM) $GA_3$ concentrations. Values are means ± SEM.

| **Genotype** | **$LER_{max}$ (mm $d^{-1}$)** | | |
|---|---|---|---|
| | **Control** | **10 uM $GA_3$** | **10 mM $GA_3$** |
| Himalaya | 28.3 ± 6.0 | 58.4 ± 1.2 | 62.7 ± 1.4 |
| *gse1a* | 9.7 ± 0.4 | 22.0 ± 0.6 | 42.1 ± 0.9 |
| *sdw3a* | 18.7 ± 0.5 | 19.5 ± 0.3 | 19.9 ± 0.3 |

**Table S4**. Maximum leaf extension rates ($LER_{max}$) in Himalaya, *grd2b, sdw3a* and *sdw3b* genotypes in response to 10 μM PBZ and 10 μM PBZ + 10 μM $GA_3$ treatments. Values are means ± SEM.

| Genotype | $LER_{max}$ (mm $d^{-1}$) | | |
|---|---|---|---|
| | **Control** | **PBZ** | **PBZ + $GA_3$** |
| Himalaya | 48.3 ± 1.1 | 25.2 ± 1.4 | 60.9 ± 0.9 |
| *grd2b* | 26.9 ± 2.2 | 19.5 ± 0.7 | 56.3 ± 2.3 |
| *sdw3a* | 24.1 ± 0.9 | 13.6 ± 1.1 | 22.5 ± 0.7 |
| *sdw3b* | 30.6 ± 1.7 | 9.3 ± 2.3 | 23.9 ± 1.9 |

**Table S5**. L1 leaf sheath length (mm) in Cadenza, *IDD5*-NS, *idd5* and *Rht-D1b* genotypes. Values are means ± SD. Data were analysed with one-way ANOVA and Tukey's post-hoc HSD test. Different letters indicate significant differences between genotypes at the 0.05 confidence level.

| Genotype | L1 leaf sheath length (mm) |
|---|---|
| Cadenza | 62.5 ± 3.8 a |
| *IDD5*-NS | 62.4 ± 6.4 a |
| *idd5* | 42.1 ± 4.7 b |
| *Rht-D1b* | 40.0 ± 7.8 b |
| *P*-value (genotype) | <0.001 |
| LSD at 5% (mm) | 3.37 |

**Table S6.** GA dose-response assays for L1 leaf sheath tissues in Cadenza, IDD5-NS, *idd5* and *Rht-D1b* genotypes. Values are means. Data were analysed with one-way ANOVA and Tukey's post-hoc HSD test. Different letters indicate significant differences between genotypes at the 0.05 confidence level.

| **Genotype** | **$H_2O$** | **$10^{-9}$ M $GA_3$** | **$10^{-8}$ M $GA_3$** | **$10^{-7}$ M $GA_3$** | **$10^{-6}$ M $GA_3$** | **$10^{-5}$ M $GA_3$** | **$10^{-4}$ M $GA_3$** |
|---|---|---|---|---|---|---|---|
| Cadenza | 62.5 a | 63.6 a | 67.0 a | 82.8 a | 103.5 a | 111.2 a | 106.1 a |
| *IDD5*-NS | 62.4 a | 61.4 a | 65.9 a | 81.7 a | 99.8 a | 107.8 a | 103.5 a |
| *idd5* | 42.0 b | 41.7 b | 43.1 b | 44.1 b | 44.3 b | 47.1 b | 45.7 b |
| *Rht-D1b* | 40.0 b | 37.3 c | 42.6 b | 44.7 b | 43.6 b | 48.1 b | 40.8 b |
| *P*-value (genotype) | <0.001 | <0.001 | <0.001 | <0.001 | <0.001 | <0.001 | <0.001 |
| SED | 1.694 | 1.552 | 1.669 | 1.982 | 2.061 | 2.271 | 2.325 |
| LSD 5% (mm) | 3.365 | 3.085 | 3.317 | 3.938 | 4.096 | 4.515 | 4.618 |

**Table S7.** GA dose-response assays for L1 leaf blade tissues in Cadenza, IDD5-NS, *idd5* and *Rht-D1b* genotypes. Data were analysed with one-way ANOVA and Tukey's post-hoc HSD test. Different letters indicate significant differences between genotypes at the 0.05 confidence level.

| **Genotype** | **$H_2O$** | **$10^{-9}$ M GA** | **$10^{-8}$ M GA** | **$10^{-7}$ M GA** | **$10^{-6}$ M GA** | **$10^{-5}$ M GA** | **$10^{-4}$ M GA** |
|---|---|---|---|---|---|---|---|
| Cadenza | 172.8 a | 171.9 a | 179.7 a | 189.1 a | 212.0 a | 213.8 a | 221.9 a |
| *IDD5*-NS | 170.2 a | 172.9 a | 183.6 a | 191.0 a | 214.0 a | 216.6 a | 224.7 a |
| *idd5* | 116.2 b | 115.9 b | 115.9 b | 121.1 b | 121.0 b | 116.5 b | 121.1 b |
| *Rht-D1b* | 136.6 c | 135.1 c | 134.9 c | 136.8 c | 147.4 c | 147.7 c | 144.6 c |
| *P*-value (genotype) | <0.001 | <0.001 | <0.001 | <0.001 | <0.001 | <0.001 | <0.001 |
| SED | 2.81 | 2.95 | 2.93 | 2.86 | 3.13 | 3.06 | 3.47 |
| LSD 5% (mm) | 5.59 | 5.87 | 5.82 | 5.69 | 6.22 | 6.08 | 6.92 |

**Table S8**. Abaxial cell lengths in 'Cadenza' and *idd5* genotypes in response to $GA_3$. Values are means ± SEM. Data were analysed using two-way ANOVA as a 2 x 2 factorial.

| | Abaxial cell length (µm) | |
|---|---|---|
| **Genotype** | **Control** | **+ 10 µM $GA_3$** |
| Cadenza | 1005.4 ± 13.6 | 1137.8 ± 17.1 |
| *idd5* | 822.3 ± 10.3 | 850.3 ± 11.5 |
| Genotype | $F$ (1,82.6) = 119.1, $P$ < 0.001 | |
| Treatment | $F$ (1,79.2) = 12.1, $P$ < 0.001 | |
| Genotype x treatment | $F$ (1,84.6) = 5.8, $P$ = 0.018 | |

**Table S9**. L1 leaf sheath lengths in ‘Cadenza’ and *idd5* genotypes in response to $GA_3$. Values are means ± SEM. Data were analysed using two-way ANOVA as a 2 x 2 factorial.

| Genotype | Abaxial cell length (µm) | |
|---|---|---|
| | Control | + 10 µM $GA_3$ |
| Cadenza | 1005.4 ± 13.6 | 1137.8 ± 17.1 |
| *idd5* | 822.3 ± 10.3 | 850.3 ± 11.5 |
| Genotype | $F$ (1,82.6) = 119.1, $P$ < 0.001 | |
| Treatment | $F$ (1,79.2) = 12.1, $P$ < 0.001 | |
| Genotype x treatment | $F$ (1,84.6) = 5.8, $P$ = 0.018 | |

**Table S10.** GA levels in leaf sheath tissues of 7-day-old seedlings from Cadenza, *Rht-D1b*, *IDD5*-NS and *idd5* genotypes. Values are mean GA levels (pg/mg) on a dry weight basis ± SE. ND = not detected. Data was analysed by one-way ANOVA and Tukey's post-hoc HSD test.

| | Cadenza | *Rht-D1b* | *IDD5*-NS | *idd5* | *P*-value (genotype) |
|---|---|---|---|---|---|
| $GA_{15}$ | ND | ND | ND | ND | - |
| $GA_{24}$ | ND | ND | ND | ND | - |
| $GA_{9}$ | ND | ND | ND | ND | - |
| $GA_{4}$ | ND | 0.261 ± 0.218 | ND | ND | - |
| $GA_{34}$ | 0.030 ± 0.004 | 0.025 ± 0.011 | 0.022 ± 0.005 | 0.022 ± 0.002 | 0.141 |
| $GA_{7}$ | ND | ND | ND | ND | - |
| $GA_{54}$ | ND | ND | ND | ND | - |
| $GA_{61}$ | ND | ND | ND | ND | - |
| $GA_{53}$ | 0.013 ± 0.007 | ND | 0.014 ± 0.007 | 0.008 ± 0.003 | 0.224 |
| $GA_{44}$ | 2.282 ± 0.442 | 0.632 ± 0.152 | 2.364 ± 0.308 | 0.396 ± 0.015 | 1.62E-07 |
| $GA_{19}$ | 1.079 ± 0.175 | 0.505 ± 0.145 | 0.794 ± 0.216 | 0.366 ± 0.049 | 2.73E-06 |
| $GA_{20}$ | 0.613 ± 0.037 | 0.817 ± 0.085 | 0.974 ± 0.092 | 0.477 ± 0.205 | 0.0272 |
| $GA_{1}$ | 1.027 ± 0.170 | 2.810 ± 0.413 | 0.290 ± 0.092 | 1.933 ± 0.071 | 4.14E-07 |
| $GA_{29}$ | 0.407 ± 0.064 | 0.084 ± 0.035 | 0.290 ± 0.063 | 0.147 ± 0.035 | 7.28E-05 |
| $GA_{8}$ | 3.174 ± 0.534 | 2.573 ± 0.317 | 4.120 ± 0.447 | 4.084 ± 0.385 | 0.00256 |
| $GA_{3}$ | 0.274 ± 0.087 | 0.394 ± 0.058 | 0.330 ± 0.185 | 0.305 ± 0.043 | 0.441 |
| $GA_{51}$ | ND | ND | ND | ND | - |

**Table S11.** Plant height, stem length, spike length and spikelet number of wild-type, *rht*-1, *idd5* and *rht-1/idd5* mutants grown in glasshouse conditions. Values are mean ± SEM. Data were analysed with one-way ANOVA and Tukey's post-hoc HSD test. Different letters indicate significant differences between genotypes at the 0.05 confidence level.

| Genotype | Plant height (mm) | Stem length (mm) | Spike length (mm) | Spikelet number |
|---|---|---|---|---|
| Cadenza | 749.1 ± 3.2 a | 642.1 ± 15.4 a | 107.0 ± 1.3 a | 19.5 ± 0.3 a |
| *rht-1* | 1107.4 ± 20.9 b | 953.0 ± 89.8 b | 154.4 ± 3.8 b | 14.5 ± 0.3 b |
| *idd5* | 643.6 ± 4.1 c | 542.1 ± 18.4 c | 104.4 ± 1.5 a | 18.7 ± 0.3 a |
| *rht-1/idd5* | 635.3 ± 6.0 c | 521.2 ± 22.6 c | 114.1 ± 1.8 c | 19.1 ± 0.2 a |
| *P*-value (genotype) | <0.001 | <0.001 | <0.001 | <0.001 |
| SED | 16.49 | 16.97 | 3.29 | 0.525 |
| LSD at 5% | 35.15 | 36.16 | 7.02 | 1.118 |

**Table S12.** Plant height of Himalaya wild-type, *sdw3a* and *sdw3b* genotypes in a wild-type and *sln1c* background grown in glasshouse conditions. Values are means ± SEM. Data was analysed with one-way ANOVA and Tukey's HSD post hoc test. Different letters indicate significant differences between genotypic classes at the 0.05 confidence level.

| Genotype | Height (cm) |
|---|---|
| Himalaya | 88.9 ± 1.1 a |
| *sdw3b* | 79.6 ± 1.7 b |
| *sdw3b/sln1c* | 93.0 ± 1.5 a |
| *sdw3a* | 66.7 ± 1.4 c |
| *sdw3a/sln1c* | 69.1 ± 1.8 c |
| *P*-value (genotype) | <0.001 |
| SED | 2.173 |
| LSD at 5% | 4.359 |

**Table S13.** Plant height of Himalaya wild-type, *sdw3a*, *sdw3b* and *sdw3e* genotypes in glasshouse conditions. Values are means ± SE. Data was analysed with one-way ANOVA and Tukey's HSD post hoc test. Different letters indicate significant differences between genotypic classes at the 0.05 confidence level.

| Genotype | Height (cm) |
|---|---|
| Himalaya | 88.7 ± 1.2 a |
| *sdw3b* | 79.6 ± 1.7 b |
| *sdw3e* | 68.6 ± 1.4 c |
| *sdw3a* | 66.7 ± 1.4 c |
| *P*-value (genotype) | <0.001 |
| SED | 2.022 |
| LSD at 5% | 4.083 |

**Table S14.** Stem length phenotypes of *Rht-D1b* and *idd5* genotypes with their appropriate control lines in glasshouse conditions. Values are means ± SE. Data were analysed using one-way ANOVA. The values significantly different between Cadenza and *IDD5*-NS are denoted by "*" and values that are significantly different between *idd5* and *Rht-D1b* are denoted by "^".

| Genotype | Stem length (mm) | Peduncle length (mm) | P-1 length (mm) | P-2 length (mm) | P-3 length (mm) |
|---|---|---|---|---|---|
| Cadenza | 686.3 ± 35.6 | 330.2 ± 46.0 | 171.8 ± 16.4 | 112.0 ± 13.5 | 62.4 ± 24.3 |
| *IDD5*-NS | 693.8 ± 38.6 | 344.6 ± 34.5 | 173.3 ± 13.6 | 113.8 ± 12.1 | 63.4 ± 21.6 |
| *idd5* | 544.6 ± 29.1***^** | 275.6 ± 30.8***** | 145.2 ± 14.7***^** | 81.4 ± 11.9***^** | 40.7 ± 15.3***^** |
| *Rht-D1b* | 508.5 ± 22.2***** | 287.0 ± 24.8***** | 133.2 ± 12.7***** | 66.2 ± 14.2***** | 22.0 ± 14.6***** |
| *P*-value (df = 69) | <0.001 | <0.001 | <0.001 | <0.001 | <0.001 |
| LSD at 5% [mm] | 15.7 | 19.5 | 8 | 7.3 | 11.4 |

**Table S15.** Height and spikelet number phenotypes of *Rht-D1b* and *idd5* genotypes with their corresponding control lines in field conditions. Values are means ± SE. Data were analysed with linear mixed models fitted using restricted maximum likelihood. Different letters indicate significant differences between genotypes at the 0.05 confidence level.

| Genotype | Plant height (mm) | Total stem length (mm) | Spike length (mm) | Peduncle length (mm) | P-1 length (mm) | P-2 length (mm) | P-3 length (mm) | Grain area ($mm^2$) | TGW (g) |
|---|---|---|---|---|---|---|---|---|---|
| Cadenza | 678.8 ± 12.0 a | 598.9 ± 11.2 a | 80.6 ± 1.6 a | 329.8 ± 6.0 a | 177.1 ± 3.9 a | 77.7 ± 4.0 a | 26.5 ± 2.9 a | 18.5 ± 0.2 a | 40.0 ± 0.5 a |
| *Rht-D1b* | 561.8 ± 14.0 b | 480.9 ± 13.2 b | 79.3 ± 1.9 a | 266.0 ± 6.9 b | 137.6 ± 5.0 b | 65.4 ± 5.0 a | 16.4 ± 3.1 b | 17.8 ± 0.3 ab | 34.2 ± 0.7 b |
| *IDD5*-NS | 682.7 ± 13.9 a | 602.3 ± 13.2 a | 80.0 ± 1.9 a | 367.8 ± 6.9 c | 153.6 ± 5.0 c | 73.2 ± 5.0 a | 15.5 ± 3.1 b | 17.6 ± 0.3 b | 36.4 ± 0.7 c |
| *idd5* | 570.3 ± 14.0 b | 491.2 ± 13.2 b | 79.0 ± 1.9 a | 256.6 ± 6.9 b | 150.2 ± 5.0 c | 70.0 ± 5.0 a | 18.1 ± 3.1 b | 16.9 ± 0.3 b | 32.4 ± 0.7 d |
| *P*-value (genotype) | <0.001 | <0.001 | 0.751 | <0.001 | <0.001 | 0.326 | 0.024 | <0.001 | <0.001 |
| SED | 15.1 | 14.6 | 2.1 | 6.8 | 6.0 | 6.0 | 3.3 | 0.3 | 0.8 |

**Table S16.** SDW3 proteins in 44 barley wheat accessions. The presence of the 15 amino acid insertion described in line Hv287 is indicated.

| Accession | SDW3 protein ID | 15 amino acid insertion |
|---|---|---|
| 10tj18 | HORVU.10tj18.PROJ.2HG00087530.1 | |
| Aizu | HORVU.AIZU_6.PROJ.2HG00087570.1 | |
| Akashinriki | HORVU.AKASHINRIKI.PROJ.2HG00119510.1 | |
| B1k-17-07 | HORVU.B1K-17-07.PROJ.2HG00084530.1 | Yes |
| B1k-33-13 | HORVU.B1K-33-13.PROJ.2HG00084920.1 | |
| Barke | HORVU.BARKE.PROJ.2HG00120340.1 | |
| Bonus | HORVU.BONUS.PROJ.2HG00088590.1 | |
| Bowman | HORVU.BOWMAN.PROJ.2HG00088620.1 | |
| F2327 | HORVU.F2327.PROJ.2HG00084260.1 | Yes |
| Foma | HORVU.FOMA.PROJ.2HG00089360.1 | |
| HID12xxi | HORVU.HID12XXI.PROJ.2HG00084220.1 | |
| HID251 | HORVU.HID251.PROJ.2HG00085320.1 | |
| HID84 | HORVU.HID84.PROJ.2HG00084180.1 | |
| HOR_10350 | HORVU.HOR_10350.PROJ.2HG00114470.1 | Yes |
| HOR_1168 | HORVU.HOR_1168.PROJ.2HG00087760.1 | Yes |
| HOR_12184 | HORVU.HOR_12184.PROJ.2HG00086820.1 | |
| HOR_13594 | HORVU.HOR_13594.PROJ.2HG00087140.1 | Yes |
| HOR_13663 | HORVU.HOR_13663.PROJ.2HG00088400.1 | |
| HOR_13942 | HORVU.HOR_13942.PROJ.2HG00117300.1 | Yes |
| HOR_14061 | HORVU.HOR_14061.PROJ.2HG00088400.1 | Yes |
| HOR_14121 | HORVU.HOR_14121.PROJ.2HG00087960.1 | Yes |
| HOR_1702 | HORVU.HOR_1702.PROJ.2HG00087910.1 | Yes |
| HOR_18321 | HORVU.HOR_18321.PROJ.2HG00089990.1 | Yes |
| HOR_21322 | HORVU.HOR_21322.PROJ.2HG00088160.1 | Yes |
| HOR_3365 | HORVU.HOR_3365.PROJ.2HG00117430.1 | Yes |
| HOR_4224 | HORVU.HOR_4224.PROJ.2HG00087910.1 | |
| HOR_495 | HORVU.HOR_495.PROJ.2HG00086980.1 | |
| HOR_6220 | HORVU.HOR_6220.PROJ.2HG00088220.1 | Yes |
| HOR_7385 | HORVU.HOR_7385.PROJ.2HG00086820.1 | |
| HOR_7552 | HORVU.HOR_7552.PROJ.2HG00118970.1 | |
| HOR_9043 | HORVU.HOR_9043.PROJ.2HG00118950.1 | Yes |
| Chikurin Ibaraki | HORVU.CHIKURIN_IBARAKI.PROJ.2HG00088340.1 | |
| Igri | HORVU.IGRI.PROJ.2HG00117850.1 | |
| Golden Melon | HORVU.GOLDEN_MELON.PROJ.2HG00087590.1 | |
| Morex | HORVU.MOREX.PROJ.2HG00116160.1 | |
| OUN333 | HORVU.OUN333.PROJ.2HG00119950.1 | |
| RGT Planet | HORVU.RGT_PLANET.PROJ.2HG00118280.1 | |
| WBDC_103 | HORVU.WBDC103.PROJ.2HG00084050.1 | |

| | | |
|---|---|---|
| WBDC_199 | HORVU.WBDC199.PROJ.2HG00084890.1 | Yes |
| WBDC_207 | HORVU.WBDC207.PROJ.2HG00085760.1 | Yes |
| WBDC_237 | HORVU.WBDC237.PROJ.2HG00085260.1 | Yes |
| WBDC_348 | HORVU.WBDC348.PROJ.2HG00084250.1 | Yes |
| ZDM01467 | HORVU.ZDM01467.PROJ.2HG00116840.1 | |
| ZDM02064 | HORVU.ZDM02064.PROJ.2HG00118750.1 | |

**Table S17.** Phenotypic data of single, double and triple *idd5* mutant genotypes in glasshouse conditions. Wild-type alleles are denoted in upper case and mutant alleles in lower case. Values are means ± SEM. Data were analysed with one-way ANOVA and Fishers post-hoc HSD test. Different letters indicate significant differences between genotypes at the 0.05 confidence level.

| Genotype | Stem length (mm) | Peduncle length (mm) | Ear length (mm) | Spikelet number |
|---|---|---|---|---|
| Cadenza | 674.7 ± 9.9 a | 351.7 ± 9.7 c | 90.7 ± 2.0 a | 17.4 ± 0.7 ab |
| *IDD5-NS* | 651 ± 11.1 a | 325.1 ± 14.9 bc | 95.4 ± 1.8 a | 17.6 ± 0.6 ab |
| aaBBDD | 653.2 ± 5.3 a | 320.5 ± 10.5 abc | 95.6 ± 2.0 a | 18.8 ± 0.6 ab |
| AAbbDD | 643.3 ± 10.6 a | 337.1 ± 11.0bc | 91.5 ± 3.3 a | 16.2 ± 0.7 a |
| AABBdd | 653.3 ± 10.9 a | 349.0 ± 9.4 c | 98.0 ± 2.3 a | 16.8 ± 0.5 ab |
| aabbDD | 661.2 ± 6.7 a | 296.9 ± 6.2 ab | 98.2 ± 2.1 a | 19.4 ± 0.3 ab |
| aaBBdd | 645.4 ± 8.6a | 328.8 ± 12.8 bc | 98.9 ± 1.8 a | 18.3 ± 0.7 ab |
| AAbbdd | 637.5 ± 8.8a | 317.3 ± 9.8 abc | 98.8 ± 1.5 a | 18.3 ± 0.5 ab |
| *idd5* | 539.7 ± 8.7 b | 273.3 ± 9.0 a | 97.3 ± 1.9 a | 18.7 ± 0.7 b |
| *P*-value (genotype) | <.001 | <.001 | 0.109 | 0.021 |
| SED | 19.60 | 14.65 | 3.165 | 0.846 |
| LSD at 5% | 40.54 | 30.30 | 6.547 | 1.750 |

**Table S18.** Sequences of primers used in this study. CAPS markers include the expected sizes of digested amplicons to distinguish genotypes.

| Oligo name | Sequence (5' - 3') | Purpose | Details |
|---|---|---|---|
| Sdw3_F1 | GGAGAGAGAGGGAGGGAAAA | *SDW3* genotyping | |
| Sdw3_R1 | CCACGGCAGAAGTCTCATTT | | |
| Sdw3_F2 | CACATCGCGCTTTCTGTG | | |
| Sdw3_R2 | GCCGTTGGGATTGTGTTG | | |
| Sdw3_F3 | GGCACACCATCCTCTCTGTT | | |
| Sdw3_R3 | TTTTCCTACCACCACCCAAC | | |
| BC1_NGS_BS_FOR | CCATCTCATCCCTGCGTGTCTCCGACTCAGCT AAGGTAACGATCGCCGCCCAAGAAGAAGAGG | *IDD5* amplicon sequencing to assess splicing variation in *IDD5-B1* | |
| BC2_NGS_BS_FOR | CCATCTCATCCCTGCGTGTCTCCGACTCAGTA AGGAGAACGATCGCCGCCCAAGAAGAAGAGG | | |
| NGS_CR_BS_REV | CCTCTCTATGGGCAGTCGGTGATGCTCCGCA CACGAACCGGTTGGTC | | |
| IDD5-A_F | GTACACCATCATCTCTGTTCCCA | CAPS marker to detect *IDD5-A1* mutation | *Sty*I digest<br>WT = 644/237/45 bp<br>MT = 644/282 bp |
| IDD5-A_R | GCCTGCGGTGAGTTGTCG | | |
| IDD5-B_WT_FAM | GAAGGTGACCAAGTTCATGCTCTCCCCGGGACGCCAGG | KASP marker to genotype *IDD5-B1* | |
| IDD5-B_MUT_HEX | GAAGGTCGGAGTCAACGGATTCTCCCCGGGACGCCAGA | | |
| IDD5-B_CR | GCAAAACCCGAAGCACGCGG | | |
| IDD5-D_WT_FAM | GAAGGTGACCAAGTTCATGCTACACAATCCCGGTTACCCC | KASP marker to genotype *IDD5-D1* | |
| IDD5-D_MUT_HEX | GAAGGTCGGAGTCAACGGATTACACAATCCCGGTTACCCT | | |
| IDD5-D_CR | AACCGGGAATGTGTTGAGC | | |
| Rht-1F3_generic | CAGATCTGCAACGTGGTGG | CAPS marker to detect *RHT-A1* mutation | *Nar*I digest<br>WT = 246/135 bp<br>MT = 381 bp |
| Rht-A1R3 | CATTTAGCTTCTTCTTCAGAGG | | |
| Rht-1F3_generic | CAGATCTGCAACGTGGTGG | CAPS marker to detect *RHT-D1* mutation | *Tsp*45I digest<br>WT = 451 bp<br>MT = 386bp/65 bp |
| Rht-D1R3 | CGGAACCACCCGTAGCCCGA | | |
